# MADRe: Strain-Level Metagenomic Classification Through Assembly-Driven Database Reduction

**DOI:** 10.1101/2025.05.12.653324

**Authors:** Josipa Lipovac, Mile Šikić, Riccardo Vicedomini, Krešimir Križanović

**Affiliations:** Laboratory for Bioinformatics and Computational Biology, Faculty of Electrical Engineering and Computing, University of Zagreb, Zagreb, Croatia; Laboratory of AI in Genomics, Genome Institute of Singapore, A*STAR, Singapore, Singapore; Univ Rennes, CNRS, Inria, IRISA - UMR 6074, F-35000 Rennes, France

**Keywords:** metagenomics, strain-level, metagenomic classification, database reduction

## Abstract

Strain-level metagenomic classification is essential for understanding microbial diversity and functional potential, but remains challenging, particularly in the absence of prior knowledge about the composition of the sample. In this paper we present MADRe, a modular and scalable pipeline for long-read strain-level metagenomic classification, enhanced with **M**etagenome **A**ssembly-Driven **D**atabase **Re**duction. MADRe combines long-read metagenome assembly, contig-to-reference mapping reassignment based on an expectation-maximization algorithm for database reduction, and probabilistic read mapping reassignment to achieve sensitive and precise classification. We extensively evaluated MADRe on simulated datasets, mock communities, and a real anaerobic digester sludge metagenome, demonstrating that it consistently outperforms existing tools by achieving higher precision with reduced false positives. MADRe’s design allows users to apply either the database reduction or read classification step individually. Using only the read classification step shows results on par with other tested tools. MADRe is open source and pub- licly available at https://github.com/lbcb-sci/MADRe.

## 1 Background

Metagenomics enables the study of genetic material from complex microbial communities found in environments such as human gut, soil, or marine ecosystems. It provides a comprehensive view of microbial diversity and interactions within these environments [1, 2]. A central challenge in metagenomic analysis is the accurate identification of organisms present in a sample, typically performed by comparing sequencing reads to reference genome databases [3].

A wide range of metagenomic classification tools have been developed, which can be broadly categorized into marker-based, DNA-to-protein and DNA-to-DNA approaches, as described in [4]. Marker-based tools, such as MetaPhlAn [5, 6], StrainPhlAn, mOTUs [7], and Melon [8], classify taxa using conserved, clade-specific marker genes. In addition to marker-based methods, SNV-based profilers (e.g., metaSNV [9] and InStrain [10], which combines both approaches) represent important strategies for strain detection and population tracking. However, most of these approaches are optimized for short-read data and rely on predefined marker sets or variant catalogs, which may not fully capture genomic diversity in complex or underrepresented microbial communities. DNA-to-protein tools, including Kaiju [11], DIAMOND [12], MMseqs2 [13], and MEGAN-LR [14] translate reads into amino acid sequences before aligning them to protein databases. DNA-to-DNA tools compare reads directly against genomic sequences and are commonly divided into k-mer-based and mapping-based tools [15]. K-mer-based tools such as Kraken2 [16], KrakenUniq [17], Bracken [18], Centrifuge [19], Centrifuger [20], CLARK/CLARK-S [21, 22], Ganon [23, 24], Taxor [25], and Sylph [26] are known for their speed and scalability to large databases, but often trade precision for speed. In contrast, mapping-based tools such as MetaMaps [27], PathoScope2 [28, 29], EMU [30] and MORA [31], which rely on read alignments and reassignment algorithms, offer higher precision at a greater computational cost.

Although k-mer-based tools, especially Kraken2 or Sylph, perform well at the species level, strain-level classification becomes increasingly challenging when sequences originate from closely related genomes [32]. However, resolving strain-level diversity is essential, as even closely related strains can exhibit substantial differences in gene content and function, with implications for microbial ecology, pathogenesis, and treatment outcomes [33, 34, 35, 36, 37].

While most existing tools are optimized for short reads due to their low cost and high accuracy, long-read sequencing technologies such as Oxford Nanopore and PacBio HiFi are rapidly improving. Longer read lengths provide advantages for genome assembly, structural-variant detection, and improved strain-level resolution.

Several short-read-based tools are designed for strain-level classification within a single species, such as StrainGE [38], StrainEST [39], and StrainSeeker [40], as well as the long-read-based ORI [41]. Other tools, including PanTax [4], MetaMaps [27], Centrifuge [19], Centrifuger [20], PathoScope2 [28, 29], and MORA [31], are suitable for more complex, multi-species datasets and support short- and long-read strain-level metagenomic classification. PanTax is a pangenome-based approach that, while supporting multi-species datasets, faces scalability limitations when applied to very large reference databases. MetaMaps is a mapping-based tool capable of high-resolution classification but is known to be extremely computationally demanding [15]. Centrifuge is a k-mer-based tool designed to perform strain-level classification, but like Pan- Tax, it encounters limitations when constructing indexes for very large reference databases. However, its successor, Centrifuger, introduces improved compres- sion and indexing strategies that enable efficient classification across large-scale genome databases. PathoScope2 is an older tool that is no longer maintained and cannot be reliably executed due to outdated dependencies and software in- compatibilities. Originally developed for strain-level classification of short reads, it is based on an expectation-maximization (EM) algorithm for read reassign- ment [42]. As part of the MORA study, the authors introduced a continuation of PathoScope2, referred to as AugPatho, which includes a modified version of the original algorithm adapted for use with long reads [31]. MORA extends this approach by combining the EM algorithm from Agamemnon [43] with a read reassignment strategy based on the Weapon-Target Assignment (WTA) prob- lem. According to its authors, MORA represents the current state-of-the-art in mapping-based long-read metagenomic classification.

Reference databases often contain multiple assemblies of the same strain and typically lack consistent organization. To address this, some strain-level classi- fication tools perform database pre-clustering according to Average Nucleotide Identity (ANI) scores. Previous studies have shown that there is no universal ANI threshold for defining strains [38, 39, 44, 45, 33]. Setting the threshold too high may erroneously separate assemblies of the same strain, whereas setting a threshold too low may incorrectly group different strains together.

Using large and diverse databases is important for accurate strain-level clas- sification [46], but it also makes the analysis much more demanding to run. MetaAlign [47] uses containment MinHash [48] to reduce the reference database prior to alignment, improving runtime while maintaining high species-level pre- cision. However, it is primarily designed for short reads, and strain-level reso- lution is not its main focus.

In this work, we introduce MADRe, a pipeline for long-read, strain-level metagenomic classification enhanced with Metagenome Assembly-Driven Database Reduction, consisting of two main phases: database reduction and read classi- fication. In the database reduction step, MADRe combines long-read assembly with an EM algorithm that assigns assembled contigs to one or more references, reducing the reference database.

In the second, read classification step, MADRe performs mappings-based read reassignment. It resolves ambiguous read mappings by assigning each read to the most likely reference, based on mapping scores and probabilistic support. We conducted an extensive evaluation of MADRe using simulated datasets, Zymo mock communities, and a real anaerobic digester sludge metagenome. The results demonstrate that MADRe achieves high precision and strain-level reso- lution while maintaining lower memory usage and runtime compared to existing tools. Additionally, the two steps of the MADRe pipeline can be run indepen- dently, and our results show that the read classification module (MADRe RC) alone performs competitively. MADRe’s approach enables the use of large, diverse reference databases spanning multiple taxonomic levels, making it well- suited for scenarios where no prior knowledge about the sample is available.

Using assembled contigs to detect potentially present strains, MADRe focuses on confidently represented organisms. As a result, compared to state-of-the-art tools, it significantly reduces the number of false positive identifications while maintaining high resolution strain-level classification.

## 2 Results

### 2.1 MADRe - method overview

We benchmarked the MADRe pipeline against state-of-the-art tools developed for the same purpose: handling large reference databases while enabling strain-level classification. These tools include MORA and AugPatho (PathoScope2), for which we evaluated both of its key modules, PathoID and PathoReport. In some of the experiments, we also evaluated k-mer based tool Kraken2, one of the most widely used metagenomic classification tools, which is often considered the standard for species-level classification. Although Kraken2 is capable of assigning reads at the strain level, its evaluation is complicated by the use of taxonomic identifiers (taxIDs) that may refer to either species or strain ranks. Despite its popularity, recent work has shown that Sylph achieves superior performance for species-level abundance estimation, reporting fewer false positives. However, since Sylph is primarily designed for abundance profiling rather than direct read classification, we did not include it in our benchmarking, which focuses explicitly on classification accuracy. Additionally, we included MADRe RC, a variant of MADRe that performs only the second step (*i.e.*, read classification) without prior database reduction.

We did not include MetaMaps, PanTax, or Centrifuge in our benchmarking analysis. In the case of MetaMaps, previous studies have reported crashes when attempting to build an index for the full Genome Taxonomy Database (GTDB), highlighting its scalability limitations [25, 31]. Similarly, our attempts to con- struct the same reference database for PanTax and Centrifuge, used successfully with other benchmarking tools, also failed due to crashes during the indexing process. Instead, we evaluated Centrifuger, a recent successor of Centrifuge that introduces improved compression and indexing strategies, enabling efficient clas- sification on large-scale genome databases.

By default, MADRe employs metaFlye [49] for assembling Oxford Nanopore (ONT) reads and metaMDBG [50] for assembling PacBio HiFi reads. To as- sess the effect of different assembly strategies on database reduction, we also performed additional experiments using Myloasm [51], a recently developed as- sembler showing promising performance on metagenomic datasets.

All commands used to run the benchmarking tools are available in the Sup- plementary File (Tools versions and commands).

### 2.2 Benchmarking details

We benchmarked the MADRe pipeline against state-of-the-art tools developed for the same purpose: handling large reference databases while enabling strain- level classification. These tools include MORA and AugPatho (PathoScope2), for which we evaluated both of its key modules, PathoID and PathoReport. In some of the experiments, we also evaluated k-mer based tool Kraken2, one of the most widely used metagenomic classification tools, which is often considered the standard for species-level classification. Although Kraken2 is capable of assign- ing reads at the strain level, its evaluation is complicated by the use of taxonomic identifiers (taxIDs) that may refer to either species or strain ranks. Despite its popularity, recent work has shown that Sylph achieves superior performance for species-level abundance estimation, reporting fewer false positives. However, since Sylph is primarily designed for abundance profiling rather than direct read classification, we did not include it in our benchmarking, which focuses explicitly on classification accuracy. Additionally, we included MADRe RC, a variant of MADRe that performs only the second step (*i.e.*, read classification) without prior database reduction.

We did not include MetaMaps, PanTax, or Centrifuge in our benchmarking analysis. In the case of MetaMaps, previous studies have reported crashes when attempting to build an index for the full Genome Taxonomy Database (GTDB), highlighting its scalability limitations [25, 31]. Similarly, our attempts to con- struct the same reference database for PanTax and Centrifuge, used successfully with other benchmarking tools, also failed due to crashes during the indexing process. Instead, we evaluated Centrifuger, a recent successor of Centrifuge that introduces improved compression and indexing strategies, enabling efficient clas- sification on large-scale genome databases.

By default, MADRe employs metaFlye [49] for assembling Oxford Nanopore (ONT) reads and metaMDBG [50] for assembling PacBio HiFi reads. To as- sess the effect of different assembly strategies on database reduction, we also performed additional experiments using Myloasm [51], a recently developed assembler showing promising performance on metagenomic datasets.

All commands used to run the benchmarking tools are available in the Supplementary File (Tools versions and commands).

### 2.3 Datasets

As part of the benchmarking process, we evaluated the mentioned tools on simulated metagenome datasets, Zymo mock communities, and a real anareobic digester sludge metagenome.

For medium-sized simulated datasets, we selected a smaller subset of genomes representing species commonly found in the human gut microbiome. Reference genomes were required to be labeled as “complete” or “chromosome” in NCBI, and to have a strain-level taxID distinct from their species-level taxID. This criterion ensured that Kraken2 could be included in the evaluation.

Using Badread tool [52] we simulated three different metagenomic datasets:

1. **sim small (4 strains)** – This dataset includes four different strain references: two strains from *Adlercreutzia equolifaciens* and two from *Streptococcus anginosus*, with varying relative abundances.
2. **sim medium (15 strains)** – This dataset contains 15 strain references distributed across five bacterial species (*i.e.*, *Helicobacter pylori*, *Cutibacterium acnes*, *Streptococcus intermedius*, *Streptococcus mutans*, *Lactococcus lactis*), with each species represented by three strains. At species level the abundances are different while strains of one species are equally abundant.
3. **sim expanded (30 strains)** – An extension of the sim medium dataset, incorporating 15 additional strains from distinct species and maintaining variable abundance levels across species. Newly added species are listed in Supplementary Table ST1.

Exact genome information including accession numbers, strain and species taxIDs, genome lengths, genome coverages, ANI values (calculated usin fastANI [44], and number of simulated reads can be found in the Supplementary Tables (ST1- ST4).

Although these datasets can be used to assess MADRe’s performance, they remain relatively simple and do not fully reflect the complexity of real metagenomic samples. Therefore, we expanded our benchmarking to include four additional simulated datasets originally used in the PanTax study [4] and we called them large-sized simulated datasets. Three of these datasets each contain 60 genomes coming from 30 species, simulated using ONT R9.4.1, ONT R10.4.1, and PacBio HiFi error profiles, respectively. These datasets were obtained directly from the PanTax Zenodo repository [53]. In addition, we generated a fourth, large-scale dataset comprising 1000 genomes from over 300 species, inspired by the CAMI challenge design. As simulated reads for this dataset were not available due to its size, we used the published reference genomes and expected abundances to simulate reads with the Badread tool. For these datasets, we additionally present distributions of ANI scores (calculated using fastANI), illustrating how many genome pairs exceed predefined ANI thresholds, as shown in Supplementary Table ST10.

We also tested MADRe using three Zymo mock communities: D6322 (ONT), D6331 (ONT) [54], and D6331 (PacBio HiFi) [55]. We included both D6311 ONT and D6311 PacBio HiFi datasets to demonstrate the pipeline’s capability across different sequencing technologies.

To evaluate performance on a complex real-world dataset, we analyzed an anaerobic digester sludge metagenome dataset sequenced using ONT R10.4.1. reads [56]. This real metagenome represents the type of scenario for which MADRe is designed, where a highly diverse sample is analyzed without prior knowledge of its taxonomic composition.

### 2.4 Database

To thoroughly evaluate strain-level classification and ensure sufficient taxonomic divergence for accurate strain-level detection, we used a database obtained via the Kraken2 interface by selecting the bacterial database. This database consists of 102,639 sequences, encompassing all RefSeq [57] complete bacterial genomes. The database was downloaded in December 2024. The exact command used for downloading the database is provided in the Supplementary File (Tools versions and commands).

For consistency, the same database was used across all tools and experiments.

### 2.5 Database Reduction

To assess the effectiveness of strain identification and database reduction, we evaluated the output of MADRe’s database reduction step on simulated datasets by comparing it to two baseline models. More precisely, assembled contigs were mapped to the large reference database and the identification of organisms was carried out using the following strategies:

- Baseline Model 1 (BM1): For the reduced database, each contig’s top reference genome, determined by the highest summarized harmonic mean mapping value (Section 5, Equation 2), was included without performing any reassignment steps. BM1 represents the ideal reduction level under the assumption of a perfect assembly, where each contig corresponds to a single strain reference.
- Baseline Model 2 (BM2): For the reduced database, each contig’s top three reference genomes, based on the highest summarized harmonic mean mapping values (Section 5, Equation 2), were included without performing any reassignment steps. BM2 defines an upper bound on the number of references expected in the reduced database. Since our simulated metagenomes contain at most three strains per species, we assume that at most three

strains could be collapsed into a single contig. This approach allows us to determine how many references should be retained in the reduced database to ensure that no true reference from the sample is missed.

The results, presented in Table 1, demonstrate that MADRe’s database re- duction achieves a high level of reduction while successfully retaining all ex- pected strains. Additionally, when compared to BM2, MADRe reduces the number of false positive species, further highlighting its effectiveness.

**Table 1:** Database reduction results. Comparison of MADRe’s database reduction performance with two baseline models on simulated datasets using large database containing **102,639** sequences: baseline model 1 (BM1), which includes only the top-1 mapping for each contig, and baseline model 2 (BM2), which includes the top-3 mappings.

|  | Metric | sim_small | sim_medium | sim_expanded |
| --- | --- | --- | --- | --- |
| number of genomes in dataset |  | 4 | 15 | 30 |
| **BM1** | # in reduced | 7 | 34 | 68 |
|  | # missing strains | 0 | 1 | 3 |
|  | # FP strains | 3 | 19 | 38 |
|  | # FP species | 1 | 0 | 6 |
| **BM2** | # in reduced | 14 | 107 | 234 |
|  | # missing strains | 0 | 0 | 0 |
|  | # FP strains | 10 | 92 | 204 |
|  | # FP species | 5 | 11 | 78 |
| **MADRe**<br>**database**<br>**reduction** | # in reduced | 7 | 42 | 84 |
|  | # missing strains | 0 | 0 | 0 |
|  | # FP strains | 3 | 27 | 54 |
|  | # FP species | 1 | 0 | 6 |

### 2.6 Classification of medium-sized simulated datasets

For medium-sized simulated datasets we compared the classification perfor- mance of MADRe, MADRe RC, MORA, AugPatho (in both PathoID and PathoReport modes), Kraken2, and Centrifuger. Since the exact source of each read is known, we define classification outcomes as follows: true positive (TP) if the read is classified under the expected strain (or expected cluster), true negative (TN) if the read is not classified and its Badread label is *random* or *junk*, false positive (FP) if the read is classified under the incorrect strain (or incorrect cluster), and false negative (FN) if the read is not classified but its label is different from *random* or *junk*.

We evaluated classification of simulated data with and without post-clustering.

The post-clustering method described in the Methods section groups closely re- lated strains based on read-to-reference mappings and assigns reads to clusters instead of individual strains. The same clustering approach was applied to all tools, utilizing read mappings to the full reference database. Kraken2’s clus- tering results are not included, as taxID alone does not allow for an accurate evaluation of post-clustering performance.

The classification results of medium-sized simulated datasets (sim small, sim medium and sim expanded) are presented in Figure 3 A., which shows the F1 scores for classification with and without post-clustering. The results demon- strate that both MADRe and MADRe RC outperform all other approaches. In- terestingly, on the sim small dataset, which includes differently abundant strains of the same species, Kraken2 performs slightly better than Centrifuger, while Centrifuger achieves higher scores than MORA and AugPatho. For the remain- ing datasets, Centrifuger performs better than Kraken2 but worse than MORA and AugPatho. In the sim expanded dataset (without clustering), MORA out- performs AugPatho’s modes, but in all other cases, including all clustering sce- narios, both AugPatho’s modes perform significantly better than MORA. When post-clustering is applied, Centrifuger shows the lowest performance across all datasets, while post-clustering further improves AugPatho’s results, bringing them close to MADRe RC. Overall, MADRe RC achieves performance compa- rable to MADRe, although this difference becomes more pronounced on more complex datasets. Figure 3B. shows the number of organisms identified by dif- ferent tools on simulated datasets. An organism is considered identified if at least one read is classified under it. In all cases, there were no false negatives — all tools successfully identified the expected organisms. However, the number of additional (false positive) identifications varies. MADRe consistently reports significantly fewer false positives. For example, in the sim small dataset, only 6 organisms were reported compared to the 4 expected, with 2 of them being extremely similar strains to those actually present in the sample. Additional metrics for organism-level identification, including TPs, FPs, FNs, accuracy, precision, recall, and F1 scores, are provided in Supplementary Table ST8, while classified read counts for each organism are listed in Supplementary Tables (ST5–ST7).

**Figure 1:**
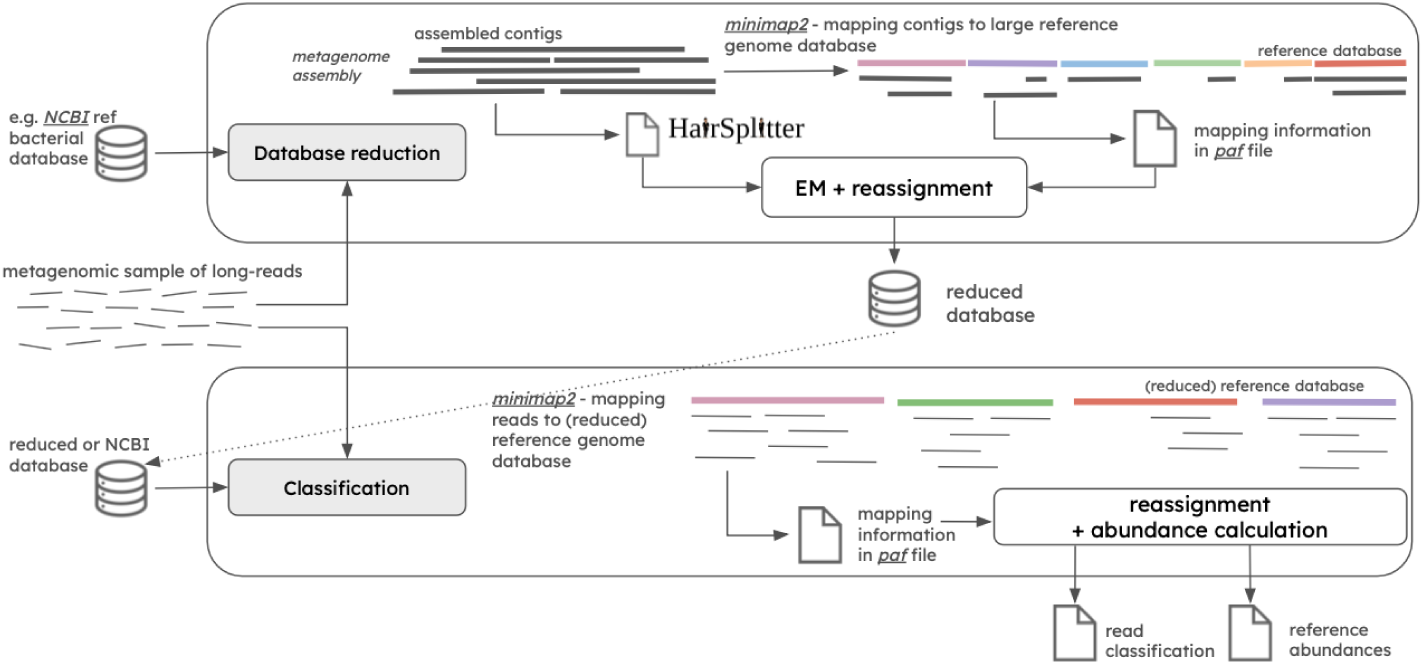
MADRe overall pipeline The first step of the pipeline performs database reduction using an EM-based contig-to-reference mapping procedure to identify organisms present in the sample. The second step involves read classification, which applies probabilistic read reassignment based on mapping information.

**Figure 2:**
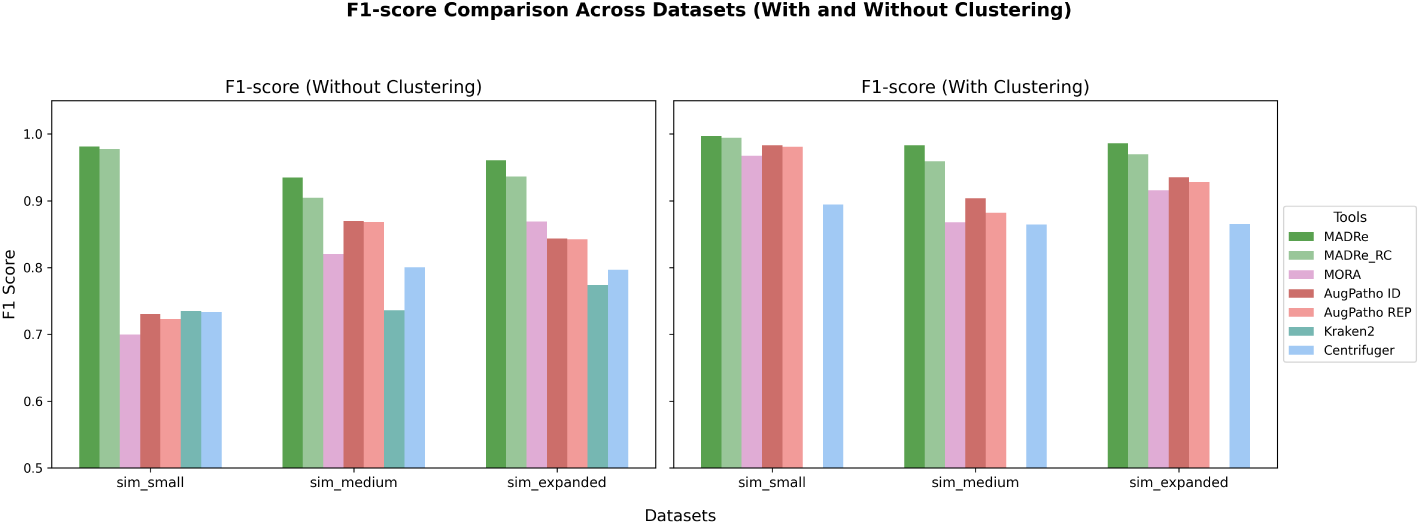
F**1 scores of strain-level classification on medium-sized simulated reads**, shown with and without post-clustering (grouping highly similar strains). Kraken2 results were omitted from the clustering analysis, as its output format does not support proper clustering.

**Figure 3:**
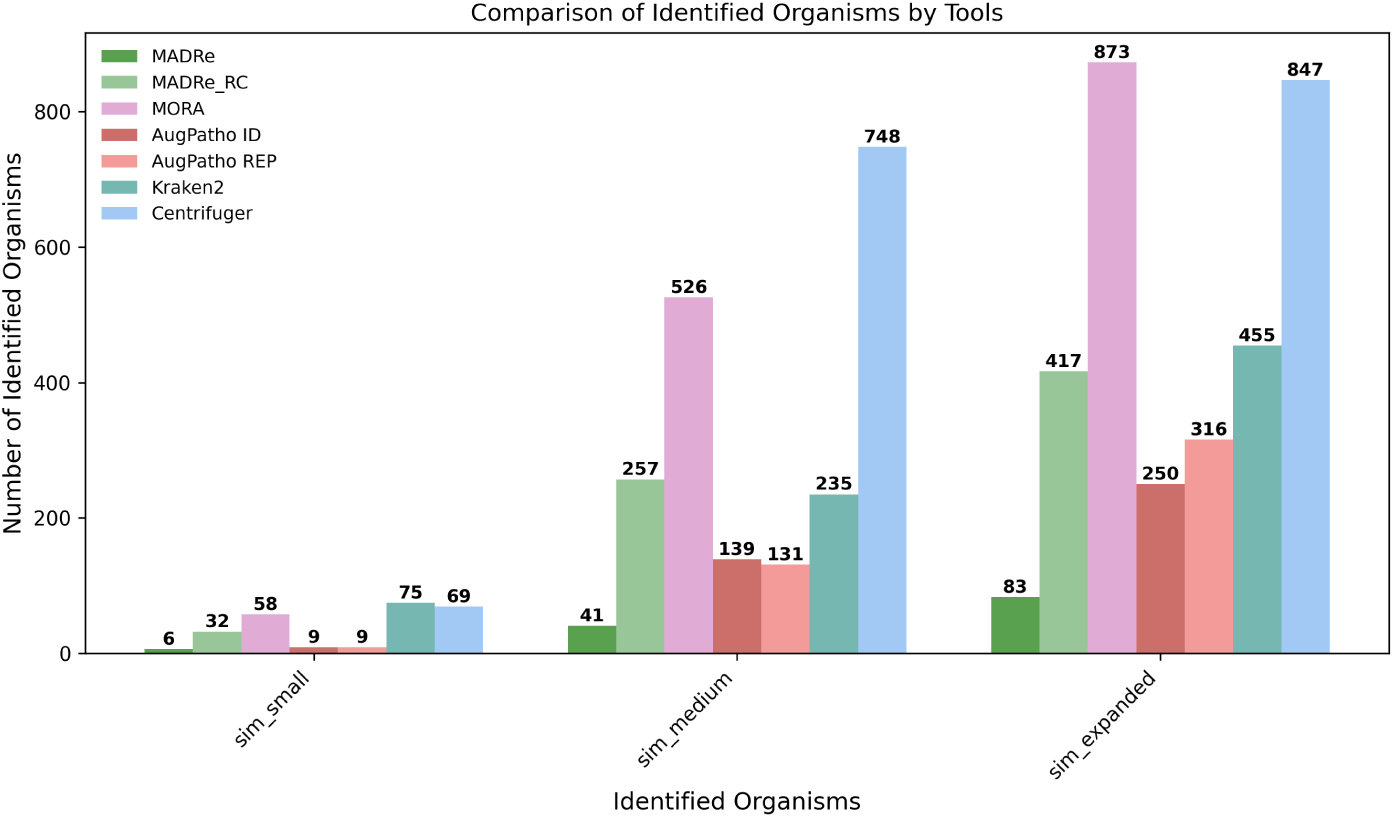
N**u**mber **of identified organisms by different tools on simulated reads.** An organism is considered identified if at least one read is classified under it. The sim small dataset contains 4 strains, sim medium contains 15 strains, and sim expanded contains 30 strains. All tools successfully identified all expected strains, resulting in no false negatives.

In addition, a Supplementary Figure S4 and Supplementary Table ST9 show Bray–Curtis (BC) distances [58] between the observed read count abundances and the ground-truth abundances, offering further insight into the similarity between the predicted and true community compositions. BC distance is one of the most commonly used distances to calculate the microbial abundance differ- ences, and is described in the Methods section. From these results, it is evident that MADRe achieves the closest match to the ground truth, while AugPatho ID reports the poorest scores among strain-level tools, including Centrifuger. Kraken2, although computationally efficient, shows the weakest overall perfor- mance.

### 2.7 Classification of large-sized simulated datasets

To further assess classification performance under more realistic metagenomic conditions, we used the simulated datasets from the *PanTax* study. We used the same evaluation procedure as the one considered for the medium-sized simulated datasets. In addition to benchmarking the standard MADRe pipeline, we also evaluated a variant in which Myloasm was used as the assembler during the database reduction step, in order to examine how different assembly approaches influence MADRe’s performance. This variant was not tested on the sim low R9.4.1 dataset, as Myloasm is not suitable for reads with that error profile.

Figure 4 presents four radar plots, each corresponding to one of the four large sized datasets, and showing F1 scores for all evaluated tools, both with and without post-clustering. (Note that post-clustering values for Kraken2 are zero, since this step was not performed for that tool.) For each tool, the best obtained F1 score is indicated in parentheses beneath its name. Across most datasets, MADRe achieved the highest scores, including its versions using different assemblers. In some cases, MADRe RC slightly outperformed MADRe, particularly on the sim high dataset. Detailed evaluation statistics are provided in Supplementary Table ST15. Exact read counts obtained from classifications are listed in Supplementary Tables (ST11–ST14), while Bray–Curtis (BC) distances between observed and expected read-count abundances are shown in Supplementary Figure S5 and Supplementary Table ST16.

**Figure 4:**
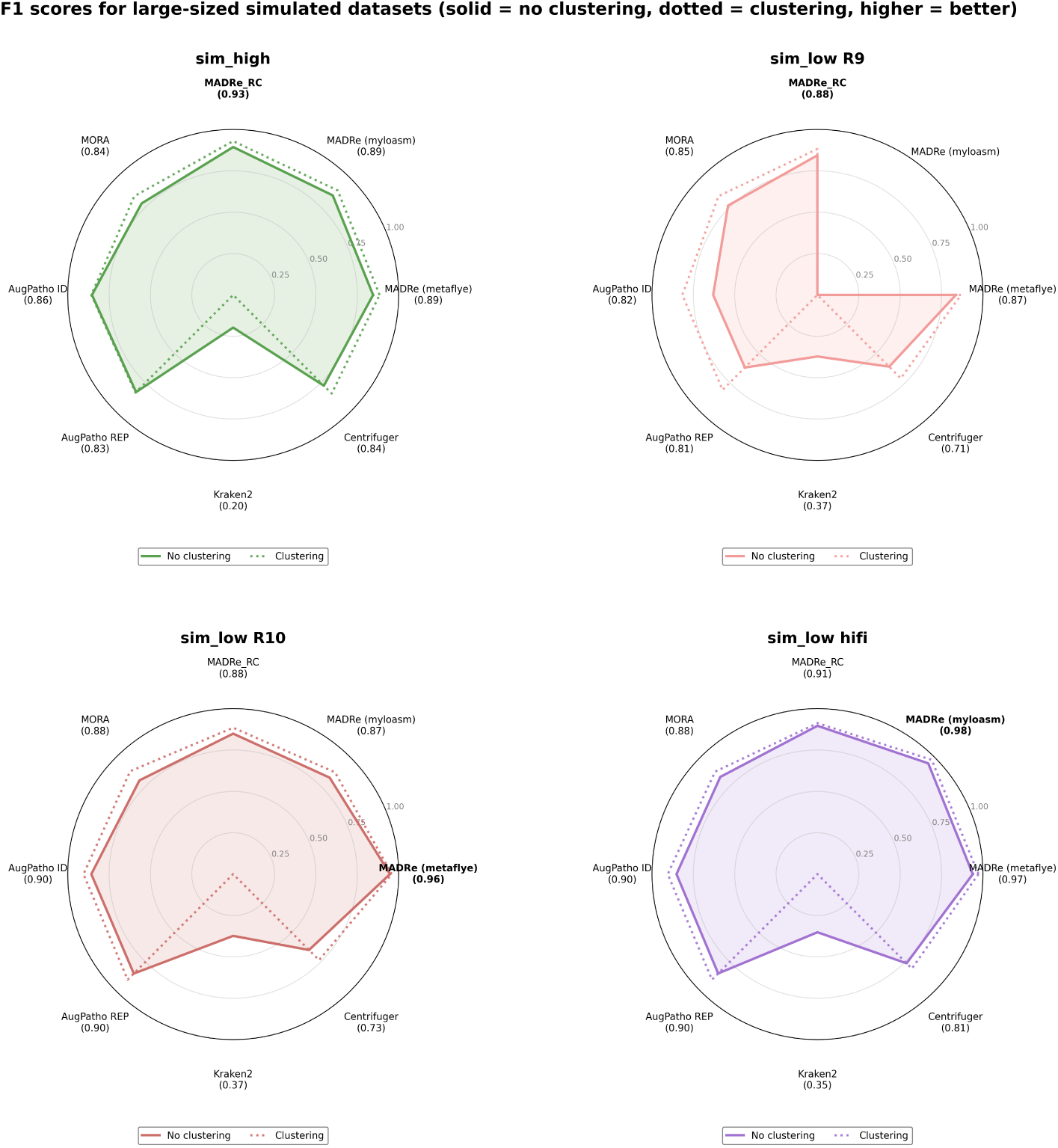
F**1 scores for large-sized simulated datasets.** Solid lines represent results without clustering, while dotted lines indicate results with clustering. Kraken2 clustering results are omitted, as its output format does not support clustering. Similarly, MADRe (Myloasm) results are excluded for ONT R9 data, since Myloasm is not designed for this type of sequencing data. In each plot, the best-performing tool is highlighted in bold, and the values in parentheses indicate the best performance achieved by each tool, with and without clustering.

Inspection of read-count abundances in the sim high dataset revealed that several genomes were not detected during the database reduction step, leading to a modest decrease in MADRe’s performance compared to MADRe RC.

When comparing assembler performance, MADRe runs based on Myloasm assemblies achieved slightly higher F1 scores than those using metaFlye. A closer look at read-count abundances revealed that 24 genomes were detected exclusively with Myloasm and 11 exclusively with metaFlye. Among the 11 missed by Myloasm, 9 belonged to the more abundant half of the community, whereas among the 24 detected only by Myloasm, just one was highly abun- dant. This pattern suggests that Myloasm-based contigs perform better for low-abundance strains, while metaFlye contigs remain more reliable for highly abundant ones.

Another observation from the sim high dataset is that, for several highly abundant strains, most benchmarking tools reported substantially lower read- count abundances than the ground truth. For example, reads originating from NZ CP012672.1 were predominantly assigned to NZ CP102233.1, a nearly iden- tical genome (ANI = 99.998) annotated under a different species (Sorangium cellulosum vs. Sorangium sp. So ce836). Because clustering was performed only among strains within the same species, this near-duplicate across species boundaries could not be resolved even after clustering.

As shown in Figure 4, all tools achieved higher F1 scores on ONT R10 and PacBio HiFi datasets compared to ONT R9, reflecting the higher base accuracy of these sequencing platforms. Interestingly, for the ONT R10 dataset, MADRe using Myloasm performed slightly worse than the metaFlye version, whereas for PacBio HiFi reads, Myloasm yielded marginally better results than the metaMDBG-based variant.

Taken together, these results, including the analyses of BC distances, demon- strate that MADRe, in all assembler configurations, consistently outperforms the other evaluated tools on the large sized simulated datasets.

### 2.8 Experiment 3 - classification of Zymo mock commu- nities reads

To evaluate MADRe on real sequencing data, we conducted experiments on three different Zymo mock community datasets: ONT Zymo D6322, which con- sists of eight organisms (seven bacterial species and one fungus), and both the ONT and HiFi versions of Zymo D6331, which contain 21 organisms, including two fungi and five different strains of *Escherichia coli*. The primary challenge in the Zymo D6331 dataset is the ability to distinguish between these closely related *E. coli* strains. In this analysis, we excluded fungal genomes, focusing solely on bacterial classifications.

For Zymo mock communities, exact reference genomes of the strains present in the sample are available, along with their theoretical relative abundances pro- vided by ZymoBIOMICS [59]. We supplemented our database with the Zymo reference genomes, assigning them separate labels. However, we did not use provided theoretical abundances in our analysis, as they may deviate from the expected values due to variations in library preparation [8, 30]. Instead, we established ground-truth read classifications. We mapped all reads to the expected bacterial reference genomes using Minimap2 and assigned true labels based on the best hit. These assignments were also used to determine the relative abundances. However, this process was not straightforward for the five

*Escherichia coli* strains, as their high similarity led to ambiguous mappings. To address this, we leveraged our clustering method (explained in *Similar strains clustering* section), which grouped these five strains into three clusters. Specifically, strains B766 and B3008 each formed separate clusters, while the remaining three strains were grouped into a single cluster, indicating that they were too similar to be reliably distinguished at the strain level. This clustering result aligns with previous findings from metagenome assembly procedures [60], where B766 and B3008 were successfully assembled, while the other three strains were not.

Benchmarking with Centrifuger and Kraken2 was not performed for this experiment, as their database construction procedures do not support the inclusion of references with custom labels, which is essential for this evaluation.

Using this information, we incorporated the clustering results into our ground-truth labeling: reads originating from the same cluster were assigned the same label, ensuring a more accurate classification.

To evaluate performance, we calculated the BC distances (eq.9) between the observed read count abundances and the ground-truth abundances, both with and without post-clustering.

Figure 5 depicts radar plots showing the BC distances for the zymo D6322 ONT, zymo D6331 ONT, and zymo D6331 HiFi datasets. Dotted lines indicate BC distances computed using only true positive classifications based on the ground truth. In the first plot, which reports results for the D6322 dataset, MADRe clearly outperforms all other tools. The second and third plots display BC distances for the D6331 ONT and HiFi datasets, respectively. For the ONT dataset, when considering all classified reads, MADRe achieves the lowest BC distance. When focusing only on true positives, MADRe and MADRe RC show comparable performance, indicating that the majority of reads classified by these tools are correctly assigned. In contrast, MORA exhibits a notably higher BC distance when evaluated only on true positives, suggesting less precise classification. For the HiFi dataset, overall distances for all the tools are significantly lower. Both AugPatho modes achieve slightly lower BC distances compared to MADRe. In Supplementary Figure S6, we present the corresponding results obtained after post-classification clustering of similar strains. Interestingly, for the ONT datasets, BC distances increased for both AugPatho and MADRe following clustering. Although the increase is not substantial, the clustering step led to elevated abundance estimates, resulting in a higher number of both false positives and true positives. This trend was not observed for the HiFi dataset, where MADRe achieved the best performance after clustering.

**Figure 5:**
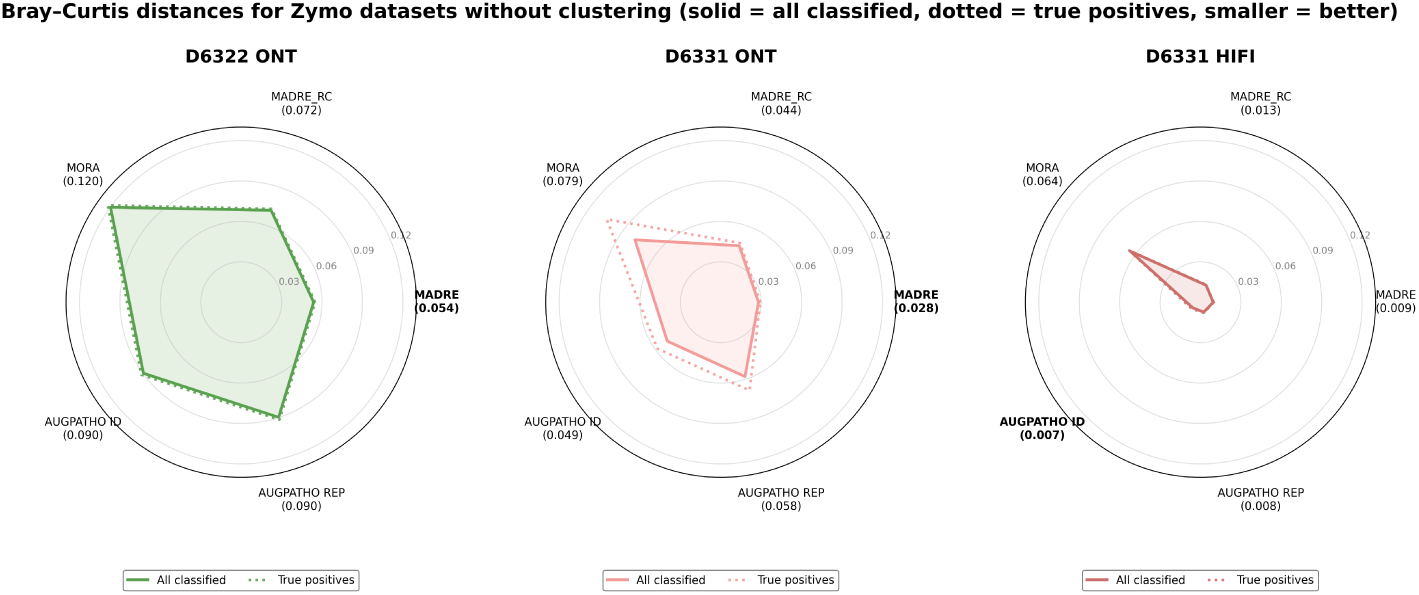
B**r**ay**–Curtis distances for Zymo datasets.** The plots show changes in BC distance with and without post-clustering of similar strains. Solid lines represent distances based on all classified read counts, while dashed lines show distances calculated using only true positive (TP) read counts.

The exact read counts, used to calculate BC distance, are listed in the Supplementary Tables (ST17-ST19).

Table 2 presents the number of false-positive species and strain identifications. MADRe reports a significantly lower number of false positives at both levels compared to other tools. Supplementary Table ST20 provides a more detailed breakdown of the number of identifications. From this table, it is evident that MADRe’s main limitation is the higher number of false negatives, primarily originating from low-abundance organisms that could not be detected using the assembly-based approach on which MADRe relies. This is further supported by the MADRe RC results, where the number of false negatives is comparable to other tools. Nevertheless, MADRe consistently reports a substantially lower number of false positives. The table also includes AugPatho results from report outputs from both modes. These reports are generated after the final reassignment step and contain only abundance estimates. Consequently, they cannot be used directly for classification evaluation. While these reports show a significantly lower number of false positives, this reduction comes at the cost of a higher number of false negatives.

**Table 2:**
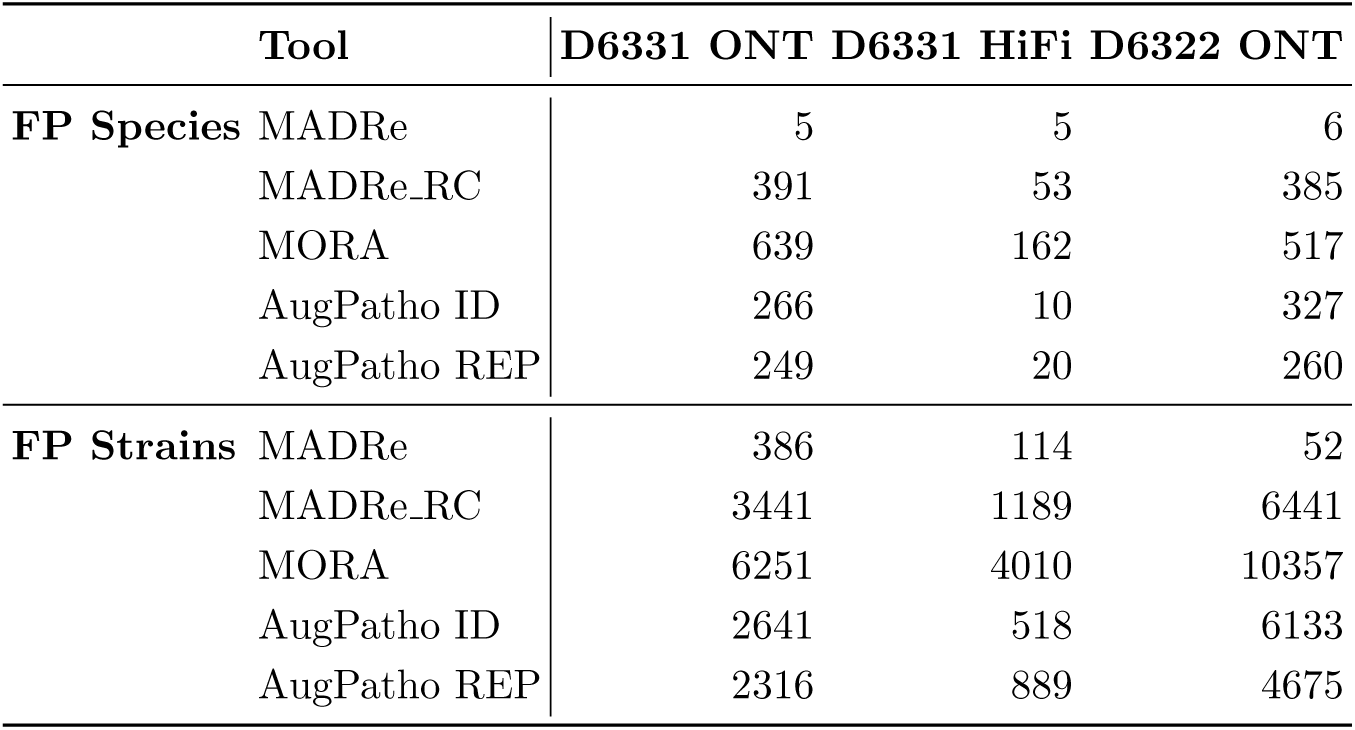
False positive (FP) species and strains detected by different tools on Zymo datasets. An organism is considered a false positive if at least one read is classified under it, but it is not in the true community.

### 2.9 Classification of real anaerobic digester sludge metagenome

While Zymo mock communities represent real metagenomic data, they do not fully capture the complexity typically found in environmental or host-associated microbial communities. To better reflect realistic classification scenarios, we evaluated MADRe and the other competing tools on a real anaerobic digester sludge metagenome. As this dataset lacks ground truth, we focused on comparative analysis of classification outputs. All results presented here include post-clustering.

In this dataset, MADRe identified 1,320 reference strains (1,502 without clustering), while MADRe RC reported 14,304 (19,067 without clustering), MORA 15,835 (23,516 without clustering), AugPatho ID 11,785 (16,488 without cluster- ing), AugPatno REP 11,134 (14,604 without clustering) and Centrifuger 23,950 (28,450 without clustering). Out of 3,646,771 total reads, MADRe classified 575,052 (∼ 16%), MADRe RC 696,961 (∼ 19%), MORA 696,839 (∼ 19%), AugPatho ID 898,906 (∼ 25%), AugPatho REP 737,714 (∼ 20%) and Centrifuger 1,350,537 (∼ 37%) reads.

Figure 6 illustrates percentile-normalized rank-abundance curves, highlighting differences in strain-level classification across the tools. The underlying read count abundance data used to generate this figure is provided in Supplementary Table ST21.

**Figure 6:**
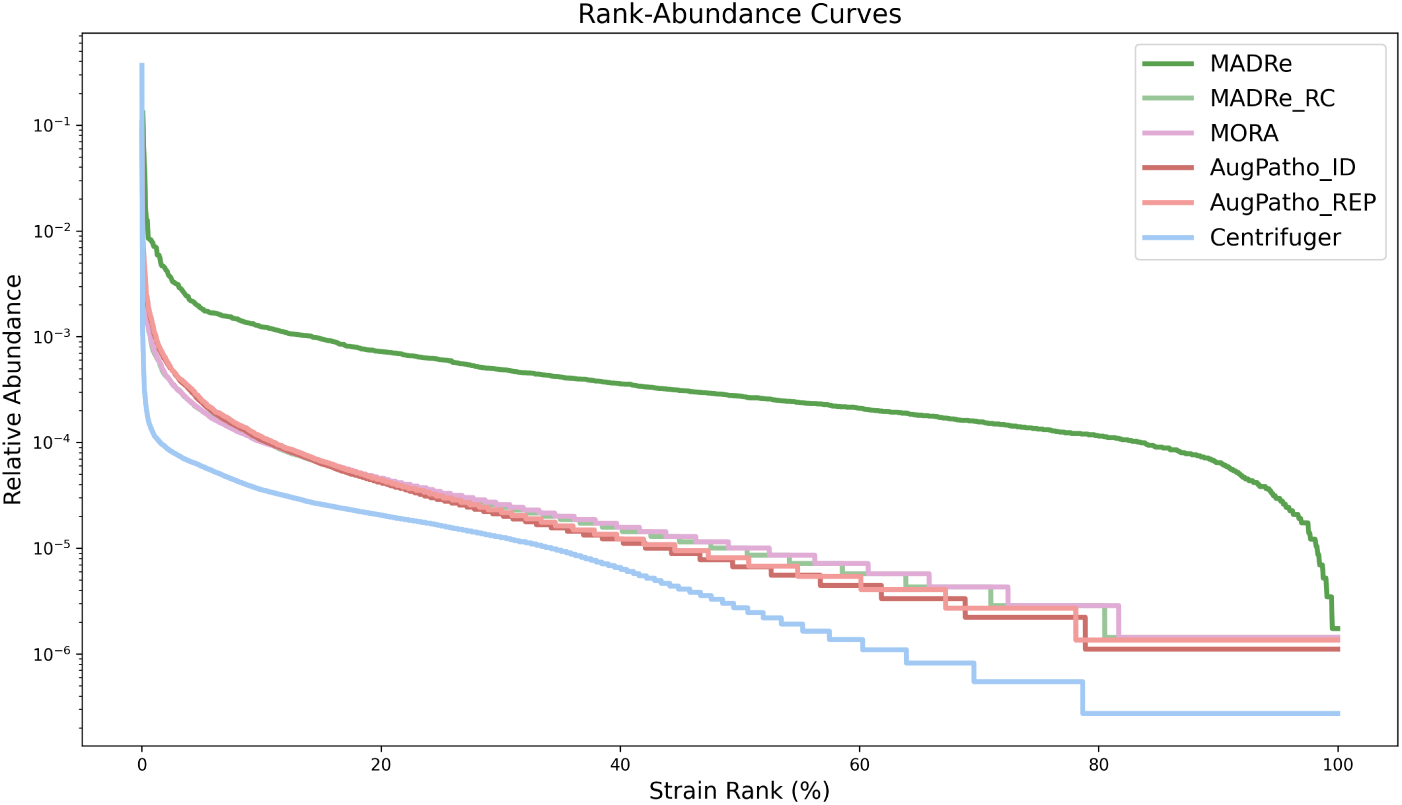
P**e**rcentile**-normalized rank-abundance curves.** The x-axis shows strain ranks expressed as percentiles, while the y-axis represents the relative abundance of each strain on a loga- rithmic scale.

The curve for MADRe displays a consistent, moderately steep gradient throughout, with notable deviations at the beginning and end. The sharp rise at the beginning indicates the presence of a highly abundant strain, significantly more dominant than the others. This can be seen for the other tools as well. Toward the end, the curve drops sharply, likely reflecting false positives or low-confidence strain assignments. Compared to the other tools, MADRe shows a smoother and more gradual decline in the abundances of lower-ranked strains. In contrast, MADRe RC, MORA, AugPatho and Centrifuger report a larger number of low-abundance strains, resulting in a more stepwise decline. The flat tail in their curves suggests that many strains are assigned near-zero abundances.

Figure 7 shows the relative abundances of the 20 most abundant strains reported by each tool, calculated relative to the total number of classified reads. We also generated an analogous visualization at the species level (Supplementary Figure S7), which additionally includes Kraken2 results. Among all tools, MADRe achieved the highest cumulative relative abundance for the top 20 strains, followed by AugPatho and MADRe RC, while MORA and Centrifuger exhibited similar but substantially lower overall contributions from their top strains. Figure 7 highlights one notable strain-level discrepancy: the strain Paludibacter propionicigenes (accession number NC 022549.1, taxID 6135), which appeared among the top 20 only in AugPatho results. To investigate this discrepancy, we examined how reads classified as taxID 6135 by AugPatho were assigned by other tools. We found that most of these reads were classified as taxID 2148 or 264636 by the other approaches. As all three of these strains belong to the *Acholeplasmataceae* family, this pattern suggests the presence of shared genomic regions and potentially an unrepresented or novel genus within this family. To further examine this case, we mapped the relevant reads to all three references and found that none yielded strong, confident alignments, indicating that the true source strain is likely missing from the reference database. We then assembled the corresponding reads into contigs and classified them using Kraken2 against the full database. In 17 contigs classified under the expected family, the highest number of k-mers matched strain 2148, although the counts were low, again supporting the hypothesis of a missing true reference. Interestingly, strain 6135 is longer than both 2148 and 264636, and prior work on MORA has shown that AugPatho’s scoring tends to favor longer, more complete genomes, which likely explains its preference for strain 6135 in this case.

**Figure 7:**
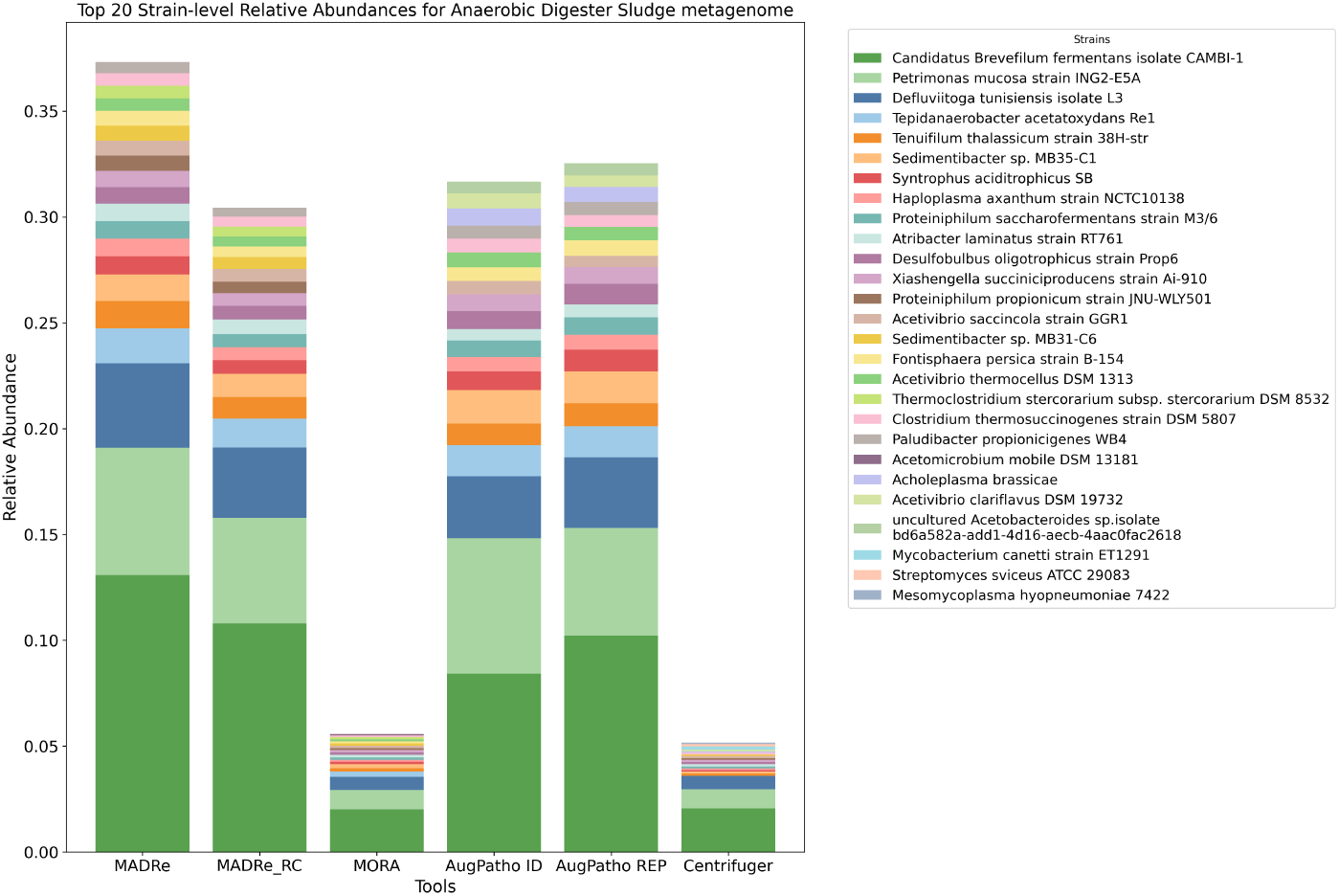
R**e**al **data strain-level abundances of the top 20 most abundant strains identified by each tool.** Two strains are highlighted in red to illustrate cases where different tools classified reads originating from an unrepresented reference to distinct false positives that share similar genomic regions.

### 2.10 Time and Memory Resources

Figure 8 presents the runtime and peak memory usage of the benchmarking tools on the ZymoD6331 ONT dataset which contained ∼1.7M reads.

**Figure 8:**
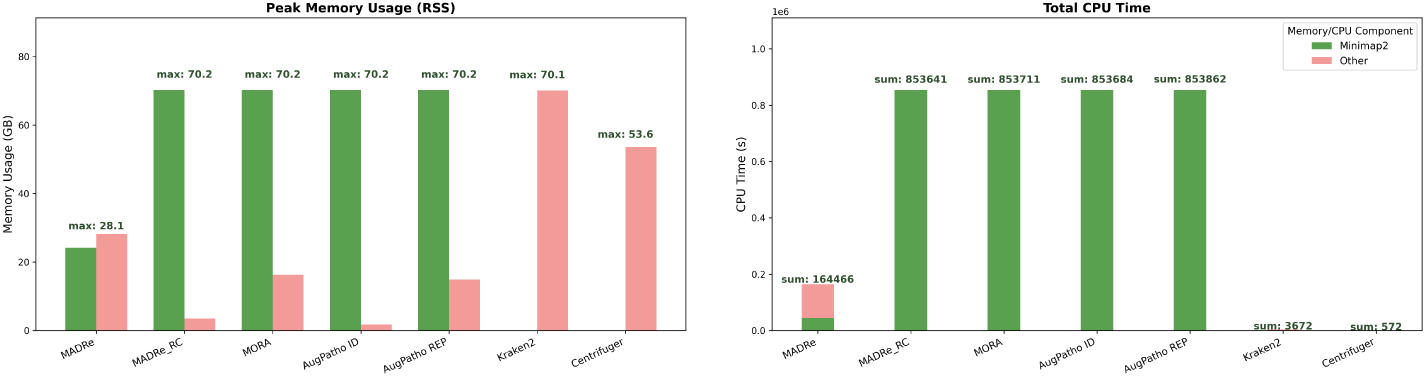
M**e**mory **(RSS peak in GB) and CPU time (in seconds)** for different tools, split between Minimap2 mapping and other processing steps.

Since majority of the tools, except Kraken2 and Centrifuger, rely on Minimap2 for read mapping, we categorized peak memory usage into components: memory used by Minimap2 and memory used by other operations. Similarly, CPU time was divided into time spent by Minimap2 and time spent on all other processing steps.

In the case of MADRe RC, MORA, and AugPatho, the “other operations” category solely consists of the read reassignment algorithm. In contrast, for MADRe it includes assembly, HairSplitter, database reduction, and read reassignment. The role of Minimap2 also differs across MADRe and other tools. In MADRe RC, MORA, and AugPatho, it is used for mapping reads to the large reference database, whereas in MADRe it is used both to map contigs to the large database and reads to the reduced one.

For HiFi reads, Minimap2 uses different parameters, and the MADRe pipeline employs metaMDBG instead of metaFlye for assembly. To account for these differences, Supplementary Table ST22 reports the same performance metrics for HiFi data.

When the dataset size increases, the situation changes. To illustrate this, we included runtime and memory usage results for the large simulated dataset sim high (containing ∼5M reads) in Supplementary Table ST22. In this case, the peak RSS for MADRe is substantially higher (exceeding 200 GB), primarily due to the assembly process, while the peak RSS for Minimap2 during read mapping to the large database remains unchanged. However, mapping reads to such a large database requires the ”–split-prefix” parameter in Minimap2, which generates temporary alignment files that are later merged at the end of the process. For this particular dataset, that procedure consumes approximately 1.2 TB of disk space, whereas the complete MADRe pipeline requires 160 GB (excluding database and read file sizes in both cases). Moreover, the entire MADRe pipeline is approximately 3.2x faster than the combination of Minimap2 with MORA or AugPatho. In contrast, Kraken2 and Centrifuge are substantially faster than mapping-based approaches and require considerably less disk space.

## 3 Discussion

In this work, we introduced MADRe, a metagenomic classification pipeline based on assembly-driven database reduction followed by read classification through mapping and reassignment. This approach enables accurate strain-level classification from large, multi-species databases without requiring prior knowledge of sample composition.

The first phase of MADRe reduces the reference database by identifying candidate strains likely to be present in the sample. Using assembly and an ex- pectation–maximization (EM) soft clustering algorithm, this step aims to retain only the relevant references while eliminating unrelated ones. To take advantage of the longer contigs produced by standard assemblers, we avoided using strain-aware metagenome assemblers such as Strainberry [61], MetaBooster [62], HyLight [62], Strainy [63], and HairSplitter [64], which are known to yield shorter contigs. Instead, we used HairSplitter’s functionality to estimate the number of collapsed strains for each contig and integrated this information with the mapping data of the initially strain-collapsed contigs. Our evaluation demonstrates that MADRe achieves effective database reduction while maintaining high recall.

Most existing strain-level classifiers either require single-species input or do not scale to large reference databases. Tools such as Kraken2, Sylph, or Centrifuger perform well at the species level and can handle large databases, making them valuable for pre-classification in strain-level workflows. However, such approaches generally require additional database preparation steps that are computationally intensive and impractical for complex metagenomic samples.

Mapping-based methods such as MORA and AugPatho represent another way to perform strain-level analysis on large databases. Nevertheless, our experiments showed that although MADRe incorporates an assembly step, typically considered both memory- and time-intensive, it required less memory than mapping raw reads directly to a large database, and even less than running Kraken2 on the same reference, when applied to a dataset of approximately 1.6 million ONT reads. For larger datasets exceeding 5 million reads, the assembly process becomes more memory demanding. However, MADRe remains substantially faster and requires considerably less disk space than mapping-based approaches. While the runtime and resource usage of Minimap2 could be reduced by using a smaller reference database, this would again require prior knowledge of the sample composition or risk omitting relevant strains. Among the evaluated strain-aware tools, MADRe is the fastest, providing an effective balance between computational efficiency and classification accuracy, and is thus well suited for scalable strain-level metagenomic analyses.

Our benchmarking analysis compared MADRe to MADRe RC, MORA, the two AugPatho modes (PathoID and PathoREP), and, in several cases, to Centrifuger and Kraken2. For AugPatho, we used the updated SAM files generated during its reassignment step, in which individual reads in some cases can be associated with multiple references. This format may improve the detection of expected references but can also introduce ambiguity, potentially contributing to higher false-positive rates. On the new simulated datasets, MADRe achieved up to a 28% improvement over other state-of-the-art methods when no clustering of similar strains was applied, and up to a 10% improvement when clustering was used. A similar trend was observed for the more complex large sized simulated datasets. Although MADRe occasionally missed low-abundance strains in these datasets, it still produced more accurate classifications than competing tools. This is particularly important since MADRe focuses on precise read-level classification rather than on abundance estimation.

A major challenge in metagenomic evaluation is the scarcity of realistic benchmark datasets, which can lead to parameter overfitting across methods, often to well-known datasets such as the Zymo communities. This may explain observations like those in the D6331 ONT dataset, where MORA and AugPatho showed substantial discrepancies between BC distances calculated from all classified reads and those derived only from true positives - the distances for true positives were notably higher. In contrast, MADRe consistently achieved better results than other tools for both evaluation types, demonstrating robust classification performance.

One limitation of MADRe observed in the Zymo benchmarks is the higher number of false-negative identifications, largely stemming from low-abundance organisms that are difficult to capture through the assembly-based approach.

However, a similar effect can be seen in AugPatho’s final reports, which include only abundance estimates from the last reassignment step - these also exhibit increased false negatives. This highlights a broader issue in metagenomic classi- fication: setting thresholds for reporting low-abundance taxa inevitably trades off between reducing false positives and increasing false negatives [65]. The identification and quantification of low-abundance organisms remain challeng- ing problems. MADRe does not apply any automatic post-filtering, leaving the decision of whether to perform additional filtering or manual investigation of low-abundance taxa to the user.

A closer look at the composition of the Zymo datasets and the definitions of ground-truth labels provides additional insight into the observed differences in tool performance. We can clearly observe performance variation across the three Zymo datasets, which can be attributed to both the sequencing technol- ogy and the evaluation methodology. As expected, the D6331 HiFi dataset yielded the best results, reflecting the higher base-level accuracy of HiFi reads compared to ONT. At first glance, it may seem surprising that performance on D6322 ONT was lower than on D6331 ONT, since D6322 contains species from different genera and should, in principle, be easier to classify. The main factor explaining this discrepancy lies in how ground-truth labels were defined. For D6322, the evaluation was straightforward - each genome represented a distinct species, and thus, an exact species-level match was required for a correct clas- sification. In contrast, D6331 includes five E. coli genomes, three of which have very high sequence identity (greater than 99.3% ANI score - calculated using fastANI). When constructing the ground truth for D6331, we clustered these three genomes and considered a read originating from any of them as correctly classified if it was assigned to any genome within that cluster. This less strin- gent criterion results in higher apparent performance for D6331 compared to D6322, an effect that applies uniformly across all evaluated tools.

In the real metagenomic dataset, MADRe classified fewer strains and focused on a confident subset of dominant organisms. In contrast, MADRe RC, MORA, AugPatho, and Centrifuger reported a much larger number of low-abundance strains. While this may suggest higher sensitivity, many of these additional de- tections are likely spurious strain-level assignments, particularly in cases where the data do not support precise strain resolution. In this dataset, certain true references were absent from the database. Under these conditions, AugPatho tended to favor longer, highly similar genomes, MORA and MADRe RC dis- persed reads across multiple low-abundance strains, whereas MADRe mostly assigned reads to the reference sharing the greatest number of similar regions with the true organism.

This behavior was further investigated through a controlled experiment on a simulated dataset containing highly similar strains, described in detail in the Supplementary File (Similar Strains Experiment). This experiment demon- strated that mapping-based tools exhibit characteristic “attractor” behavior, often failing to proportionally distribute reads among near-identical strains. AugPatho and MORA, which rely on probabilistic or abundance-constrained models, frequently collapsed or misallocated reads to different representatives when the database or dataset composition changed. In contrast, MADRe con- sistently assigned reads to the most similar available reference, the centroid, thereby maintaining stable classifications. When centroid references were re- moved, MADRe dynamically adapted by reallocating reads to the closest rep- resentative. These results highlight that MADRe’s centroid-based strategy en- sures stable and interpretable performance in challenging scenarios where am- biguity among nearly identical genomes is unavoidable.

These observations also emphasize a broader limitation of current long-read metagenomic classifiers: all existing methods struggle to resolve strains at ex- tremely low sequence divergence. For this reason, in our evaluation we addi- tionally report results at the cluster level, where highly similar genomes are grouped together based on their mapping profiles. This approach avoids penal- izing tools for inevitable redistribution within such groups and provides a more biologically meaningful measure of performance. Unlike conventional clustering by average nucleotide identity (ANI), our method groups references according to shared mapping profiles, focusing on patterns reflected in the data rather than static reference similarity. This design supports the concept of sample-aware reference groupings that better capture functional and ecological relationships and could enhance classification accuracy in the presence of closely related or- ganisms. Such clustering could also guide adaptive reference construction or real-time database refinement as additional samples are analyzed. Although clustering was used only for evaluation in this study and applied uniformly across all tools, future work will include deeper investigation of this method and its integration into the full MADRe pipeline.

MADRe is a modular pipeline composed of independent components, allow- ing easy adaptation to different tools, such as alternative assemblers. In the large sized simulated read experiments, we evaluated a MADRe version that used the Myloasm assembler instead of metaFlye and metaMDBG. The overall results were comparable. However, Myloasm showed better detection of low-abundance strains, whereas metaFlye performed slightly better for highly abundant ones. These findings indicate that different assemblers may be advantageous for dif- ferent use cases. Consequently, we included Myloasm as an optional component within the MADRe pipeline, and future versions will support additional assem- blers and related tools.

Beyond strain-level classification, MADRe’s modular design, particularly its database reduction and probabilistic reassignment components, offers poten- tial for broader applications. These include contig binning, assembly refine- ment, and functional gene profiling, where confident reference reduction and ambiguity-aware read handling are equally valuable.

## 4 Conclusion

In this study, we introduced MADRe, a novel pipeline for strain-level metage- nomic classification of long-read sequencing data. MADRe combines long-read assembly, EM-based contig-to-reference mapping reassignment for database reduction, and probabilistic read reassignment to deliver accurate and efficient classification, even without prior knowledge of sample composition. Unlike many existing tools, MADRe is designed to operate with large, diverse databases spanning multiple taxonomic levels, enabling high-resolution classification while minimizing false positives.

The pipeline consists of two distinct steps: database reduction and read classification, both of which can be executed independently. If general insight into the strains present in a sample is required, the first step can be used alone. Conversely, when prior knowledge about the sample exists, or when a reduced reference set is already available, the read classification step can be applied independently. MADRe provides a practical, scalable, and modular solution for strain-level classification in complex microbial communities.

## 5 Materials and Methods

### 5.1 MADRe Database Reduction

The database reduction step, shown in Figure 1 and illustrated in more detail in Supplementary Figure S1, consists of two main phases: input file preparation and the database reduction.

In the input file preparation phase, raw long metagenomic reads are first assembled using metaFlye for ONT reads or metaMDBG for HiFi reads. When multiple strains of the same species are present in a sample, the assembly process can lead to strain collapse, producing contigs that represent a blend of closely related strains rather than distinct strain-specific sequences. Instead of using strain-aware metagenome assemblers, which typically generate shorter contigs, we chose to retain the longer contigs and infer strain-level complexity using HairSplitter functionality which estimates the number of collapsed strains per contig.

Assembled contigs are mapped to the reference database using Minimap2 with the *asm5* parameter preset, generating a PAF file as output. We chose this preset because, compared to *asm10* and *asm20*, it provides higher sensitivity, which is crucial for capturing more accurate and complete alignments of contigs to highly similar reference genomes. The MADRe database reduction process takes two key inputs: the estimated number of collapsed strains per contig determined by HairSplitter and the contig-to-reference mappings from Minimap2.

The database reduction process is based on the EM algorithm, which reassigns contigs to different references while performing soft clustering, allowing a single contig to be assigned to multiple references with different probabilities. The EM algorithm is widely used for handling ambiguous mappings in metagenomic classification [31, 29, 28, 30, 66, 43]. The implementation of the EM algorithm in MADRe is inspired by PathoScope2 [29] and EMU [30].

The database reduction process consists of three main steps. We can define a set of contigs as *C* = {*c*_1_*, c*_2_*, . . ., c_x_*}, where *x* is the number of contigs in the assembly. The set of references to which at least one contig is mapped is defined as *R* = {*r*_1_*, r*_2_*, . . ., r_g_*}, where *g* is the number of references. Additionally, let *M* represent the set of all of the mappings. In the first step, we compute a mapping score H for each mapping in the PAF file using the equation:

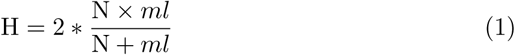

which represents the harmonic mean between the exact number of matches N and the mapping length *ml*. The *ml* is defined as the maximum value between the query mapping length and the reference mapping length. Applying the harmonic mean allows us to emphasize the smaller value, ensuring that a mapping does not receive an inflated score due, for example, to a very long but low-quality alignment.

The summarized mapping value S is then calculated for each contig-reference pair using:

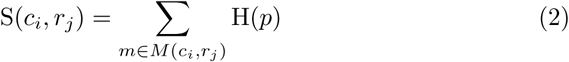

where *M* (*c_i_, r_j_*) represents the set of mappings of contig *c_i_* to reference *r_j_*. This ensures that S(*c_i_, r_j_*) is computed by summing the mapping values of all instances where *c_i_* maps to *r_j_*, thus capturing all possible alignments between the contig and the reference. Following this, we divided mappings into *unique* and *non-unique*. Unique mappings occur when a contig maps exclusively to a single reference, whereas non-unique mappings represent ambiguous cases that require further resolution.

In the second step, non-unique mappings are processed using the EM algorithm, which iteratively refines contig assignments based on mapping probabilities. The E-step updates the expected assignments of contigs, while the M-step re-estimates the parameters using the newly computed assignment probabilities from the previous iteration.

The probability of selecting a reference *r_i_* is given by:

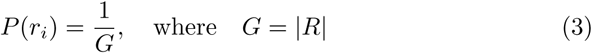

The conditional probability of *c_i_* given *r_i_* is expressed as:

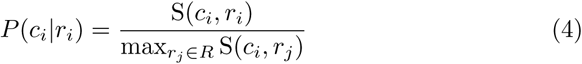

The log-likelihood function *L*(*X*) is given by:

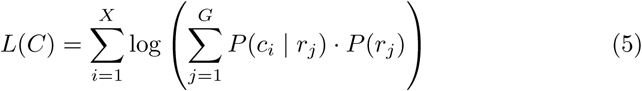

The expectation step (E-step) updates the posterior probability *P* (*r_i_* | *c_i_*) as follows:

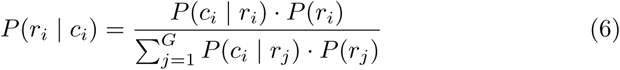

The maximization step (M-step) updates the prior probability *P* (*r_i_*) as follows:

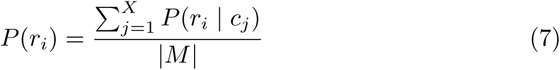

The EM algorithm runs iteratively until it converges or reaches the maximum number of iterations set by the stopping criteria. Once the algorithm outputs posterior probabilities, these values are used to determine which references will be included in the reduced database.

Before selecting references, we first classify each contig at the species level. This is done by summing the posterior probabilities across all references belong- ing to a species and assigning the contig to the species with the highest total probability. After determining the species classification, we retain only poste- rior probabilities associated with references belonging to the selected species. Finally, for each contig, we select N + 2 reference genomes to include in the re- duced database. The value of N is estimated based on the number of collapsed strains identified by HairSplitter. By default, MADRe adds two additional ref- erences to avoid excluding expected strains, although this offset can be adjusted through user parameters.

### 5.2 MADRe Read Classification

The read classification step in MADRe is designed to operate both with and without prior database reduction. The only requirement is a PAF file, the Minimap2 output containing read-to-database mappings, where each database sequence includes the corresponding taxonomic identifier. The MADRe read classification workflow is illustrated in Supplementary Figure S2.

The process begins by computing a mapping score for each alignment, de- fined as the ratio between the number of exact matches (*N*) and the mapping length (*ml*):

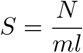

Mappings are then categorized into unique and non-unique. Since a read can have multiple alignments, only the alignment with the highest score is retained for each read-reference pair. If a read has a single best-scoring mapping, it is classified as a unique mapping. Conversely, if multiple mappings share the same highest score, they are considered non-unique, and the read will go through a reassignment process. Formally:

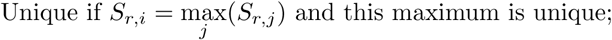

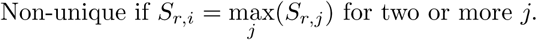

Before reassigning non-unique mappings, reads are first classified at the species level. For each read *r*, the maximum mapping score among all references belonging to a species *s* is computed as:

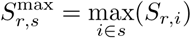

The species with the highest *S*^max^ is selected as the species-level assignment for that read. Although the default assumption is that a read can be uniquely mapped to a single species but may map ambiguously to multiple strains within that species, this assumption does not always hold. In rare cases, two genomes from different species may share highly similar regions, making it difficult to determine the true origin of a read. In such situations, reads are randomly distributed between the corresponding species. These cases are uncommon and typically arise from taxonomic inconsistencies, for example when nearly identical strains according to taxonomy belong to different species.

To reassign non-unique mappings at the strain level, a species-specific clustering algorithm is applied. This algorithm evaluates the number of unique and non-unique mappings associated with each reference within the same species, identifying groups of references that share many mappings, indicating that they likely represent overlapping genomic regions and should form clusters.

The fundamental assumption is that a reference truly present in the sample should accumulate the highest number of mappings (both unique and high-confidence non-unique). Let *M_i_* denote the total number of mappings to refer- ence *i*:

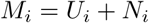

where *U_i_* and *N_i_* represent the counts of unique and non-unique mappings, respectively. The expected references are identified as those with the highest *M_i_* within each cluster, and all non-unique reads are reassigned to the most probable reference in that cluster:

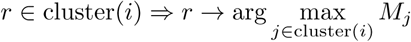

This procedure ensures that ambiguous reads are redistributed toward references that are both well supported by unique evidence and consistent with the mapping structure observed across the sample. By reassigning reads to the most representative reference within each cluster, this methodology establishes MADRe’s centroid-based behavior, maintaining stable and interpretable classifications even in the presence of highly similar strains.

#### 5.2.1 Abundances calculation

At this stage, each read is assigned to a single reference genome. Based on these assignments, MADRe calculates the abundance of each detected strain.

The primary abundance output file reports the number of reads assigned to each reference. However, MADRe also provides an option to compute a length- normalized abundance, which accounts for both read and reference lengths. This alternative abundance metric is calculated as:

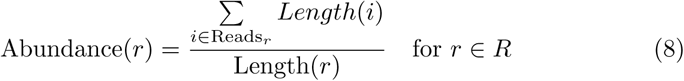

where *R* is the set of the references and *Reads_r_* is set of the reads classified under reference *r*.

#### 5.2.2 Similar strains clustering

The high similarity between closely related strains and the lack of a clear thresh- old for defining when two sequences represent the same strain, makes it difficult to ensure that a reference database contains only unique strain sequences [33]. Some entries may correspond to highly similar strains or even to multiple assem- blies of the same strain. To address this, MADRe includes an optional reference clustering step within the read classification process, designed to group similar references based on shared read mappings.

This clustering step operates on the same mapping file used in the clas- sification stage and produces two output files: one reporting the abundances of identified clusters and the other listing the representative reference for each cluster.

The clustering procedure is illustrated in Supplementary Figure S3. It begins by using the calculated mapping scores and species-level labels. For each species- specific subset, the algorithm identifies the highest-quality mapping for each read. A binary vector is constructed for each reference, where each bit indicates whether a given read strongly supports that reference. These binary vectors are then clustered using DBSCAN with precomputed Jaccard distances, eps = 0.9, and min samples = 1. Cluster-level abundances are computed accordingly. As a result, the post-clustering abundance files report only the representative references for each cluster.

### 5.3 Evaluation details

Our evaluation is primarily focused on exact read-level taxonomic assignments and read count-based abundance estimates.

To ensure a fair comparison, all tools were benchmarked using the same large reference database. For tools requiring mapping files as input, namely MORA, AugPatho, and MADRe RC, we used a unified set of read-to-reference alignments generated by Minimap2. All reads were mapped to the full database, and the resulting SAM file was used directly for MORA and AugPatho. This SAM file was subsequently converted to PAF format using Paftools, as required by MADRe RC.

Simulated reads were generated with the Badread tool [52], applying the whole-metagenome simulation mode with default values for chimeric, junk, and random reads. The simulation was performed using the *nanopore2023* model, which corresponds to the ONT R10.4.1. The exact command used is provided in the Supplementary File.

In case of simulated datasets, ground truth was available for every read, including its corresponding strain-level taxID and reference accession. Using this information, we computed true positives (TP), false positives (FP), true negatives (TN), and false negatives (FN) by comparing the assigned strain-level taxIDs with the expected ones. For Kraken2, we extracted read IDs and assigned taxIDs from its output. If a read was assigned to a higher taxonomic level, even if it was taxonomic correct, we treated it as a false positive, as the evaluation strictly focused on strain-level classification.

For the simulated datasets and Zymo mock communities we calculated Bray-Curtis (BC) distance as:

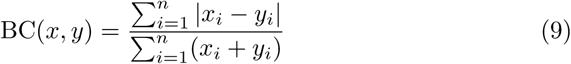

Where *x* = (*x*_1_*, x*_2_*, . . ., x_n_*) and *y* = (*y*_1_*, y*_2_*, . . ., y_n_*) are the abundance vectors for two samples or profiles, *n* is the number of strains, *x_i_* and *y_i_* are the abundances of the *i*^th^ strain in samples *x* and *y*, respectively. BC(*x, y*) is the Bray-Curtis dissimilarity or distance, ranging from 0 (identical composition) to 1 (completely disjoint).

In the case of medium-sized simulated datasets (sim small, sim medium and sim expanded) and the real dataset, the evaluation was also performed at the species level (results presented in Supplementary Table ST8 and Supplementary Figure S7). All strain-level classifications were uplifted to their corresponding species, and read count abundances were calculated. For Kraken2, we used the species-level abundances reported in its summary file, limited to entries labeled with an “S” (species rank).

MADRe, MADRe RC, and MORA each produce read-level classification files that associate each read with a reference genome. Since all tools shared the same mapping input, for AugPatho we ran PathoID and PathoREPORT steps, which output an updated SAM files and a report containing reference abundance estimates. In this updated SAM files, a single read can be associated with multiple references. For evaluation purposes, we allowed such multi-reference assignments, which may slightly benefit AugPatho by increasing the number of true positives, while also increasing the risk of false positives. These trade-offs are largely neutralized when clustering is applied, as similar strains typically end up grouped in the same cluster. With Centrifuger we encountered one limitation - Centrifuger cannot confidently assign a read to a specific reference sequence (e.g., when multiple chromosomes belong to the same strain), it often classifies the read under the NCBI strain-level taxid. In some cases, this strain taxid is identical to the species taxid, making it impossible to directly and fairly compare such classifications to those of other tools that operate at the sequence level. For benchmarking consistency, we therefore considered as true positives only the reads correctly classified under the expected reference sequence. It is important to note that this issue affected a relatively small fraction of reads (approximately 9000 out of 5 million reads in the 1000-genome dataset).

We used the same clustering across all tools to ensure consistency in cluster-based evaluation. Our clustering is based on read-to-reference mapping profiles, which can differ depending on the size and composition of the database. For example, when reads are mapped to a reduced database, the absence of certain similar references can make ambiguous mappings more resolvable. To avoid such inconsistencies, clustering was performed only once on the PAF file used for MADRe RC, which contains read mappings to the complete reference database. During cluster-level evaluation, a classification was considered a true positive if the read was assigned to a reference that belongs to the same cluster as the ground truth reference:

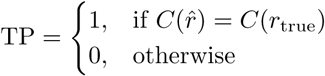

where *C*(*r*^) denotes the cluster of the predicted reference and *C*(*r*_true_) denotes the cluster of the true reference. A classification is considered a true positive (TP) if both references belong to the same cluster.

All commands used to perform classification with the evaluated tools are listed in the Supplementary File.

## 6 Data availability

The source code for MADRe is avaliable at https://github.com/lbcb-sci/MADRe. Simulated medium-sized data can be accessed via Zenodo. Zymo D6322 ONT dataset is obtained from BioProject PRJNA1240873, zymo D6331 ONT dataset if obtained from [54], and zymo D6331 PacBio HiFi dataset from [55].

## Acknowledgments

The authors thank Lune Angevin for testing the tool and providing valuable feedback.

## 7 Contributions

K.K. and M.S^̌^. conceived the study. J.L. designed and implemented the pipeline. K.K. supervised database reduction implementation. R.V. supervised read classification implementation. J.L. drafted the manuscript. K.K., R.V. and M.S^̌^. revised the manuscript. All authors read and approved the manuscript.

## 8 Funding

This work was supported by the Croatian Science Foundation under grants IP- 2018-01-5886 (SIGMA) and MOBDOK-2023-2941, and by the Singapore Ministry of Health’s National Medical Research Council, Singapore, under the grant MOH-000649-01 (Rapid diagnostic of infectious diseases based on nanopore sequencing and AI methods) – Individual Research Grant (NMRC/OFIRG/MOH- 000649-00).

## 9 Competing interests

M.S^̌^. has been jointly funded by Oxford Nanopore Technologies and AI Sin- gapore for the project AI-driven De Novo Diploid Assembler. The remaining authors declare no competing interests.

## Notes

### Competing Interest Statement

M.S. has been jointly funded by Oxford Nanopore Technologies and AI Singapore for the project AI-driven De Novo Diploid Assembler. The remaining authors declare no competing interests.

### Summary of Updates

New datasets added; Figures, tables and results revised; Supplemental files updated.

