## Supplementary File for "MADRe: Strain-Level Metagenomic Classification Through Assembly-Driven Database Reduction"

#### Tool versions and commands

##### Data simulation - **badread** - version: v0.4.1

```
$ badread simulate --reference reference_file.fasta \
--quantity 10x > reads.fastq
```

File `reference_file.fasta` contained organisms that are listed in Supplementary table ST1.

##### Database building - **Kraken2** - version: 2.1.3

```
$ kraken2-build --download-taxonomy --db $DBNAME

$ kraken2-build --download-library bacteria --db $DBNAME

$ kraken2-build --build --db $DBNAME
```

In the case of all of the tools we used the .fna file from
`$ DBNAME/library/bacteria/library.fna` as a starting main database.

##### Classification - **Kraken2** - version: 2.1.3

```
$ kraken2 --db $DBNAME --threads 64 --output k2.out --report
k2.report reads.fastq
```

##### Mapping reads (ONT and HiFi) to database - **minimap2** - version: 2.28 - **samtools** - version: 1.15.1

```
$ minimap2 -ax map-ont (map-hifi) database_file.fasta reads.fastq
-t 64 > reads_to_db.sam
$ paftools.js sam2paf reads_to_db.sam > reads_to_db.paf
```

##### Classification - **MORA** - version: 1.0

```
$ mora --sam reads_to_db.sam --out reads.mora -t 64
```

##### Classification - **AugPatho - ID module** - version 8.30

```
$ MORA-data/AugPatho2/pathoscope2.py ID -alignFile reads_to_db.sam
\
-outDir out_ID
```

#### Classification - **AugPatho - Report module** - version 8.30

```
$ MORA-data/AugPatho2/pathoscope2.py REP -samfile out_ID/ID.sam\
-outDir out_REP
```

#### Database building - **Centrifuger** - version

```
$ centrifuger-build -r reference_file.fasta -l
lisit_of_references.txt  -o centrifuger_index -t 64
```

#### Classification - **Centrifuger** - version

```
$ centrifuger -x centrifuger_index -u reads.fastq -t64 >
output.tsv
```

#### MADRe pipeline

- Metagenome assembly ONT - **metaFlye** - version: 2.9.5-b1801

```
$ flye --nano-raw ont_reads.fastq -t64 --meta --out-dir
metaflye_out
```

- Metagenome assembly HiFi - **metaMDBG** - version: 1.1

```
$ metaMDBG asm --out-dir metaMDGB_out --in-hifi reads.fastq
--threads 64
```

- Metagenome assembly - **Myloasm** - version: 0.2.0

```
$ myloasm reads.fastq -o myloasm_out -t 64 (--hifi)
```

- Mapping contigs to database - **minimap2** - version: 2.26-r1175

```
$ minimap2 -x asm5 database_file.fasta assembly.fasta -t 64 >
asm_to_db.paf
```

- Collapsed strains calculation - **HairSplitter** - version: v.1.9.18

```
$ hairsplitter.py -i assembly.fasta -f reads.fastq -t 64 -o
hairsplitter_out
```

- Database reduction:

```
$ python DatabaseReduction.py --paf_path asm_to_db.paf
--num_collapsed_strains collapsed_contigs.txt --reduced_list_txt
reduced_list.txt --reduced_db reduced_db.fasta --threads 64
```

- Mapping reads to reduced database - **minimap2** - version: 2.26-r1175

```
$ minimap2 -cx map-ont (map-hifi) reduced_db.fasta reads.fastq -t
64 > reads_to_reduced_db.paf
```

- Classification

```
$ python ReadClassification.py --paf_path reads_to_reduced_db.paf
```

- Whole pipeline - **MADRe** - version: 0.0.4

```
$ python MADRe.py --out-folder madre_out --reads reads.fastq
--reads_flag ont (hifi) --threads 64
```

#### Clustering - **MADRe**

```
$ python ReadClassification.py --paf_path reads_to_db.paf
--clustering_out clustering_dir
```

In the case of all of the tools we used the same clustering information obtained with this command and for DBSCAN's *eps* parameter we used 0.9 value.

Abundance calculation - **MADRe**

```
$ python CalculateAbundances.py --reads reads.fastq --read_class
classification.out (--clusters clustering_dir)
```

In the case of all of the tools we used the same way of abundance calculation which is based on the read count.

### Database Reduction

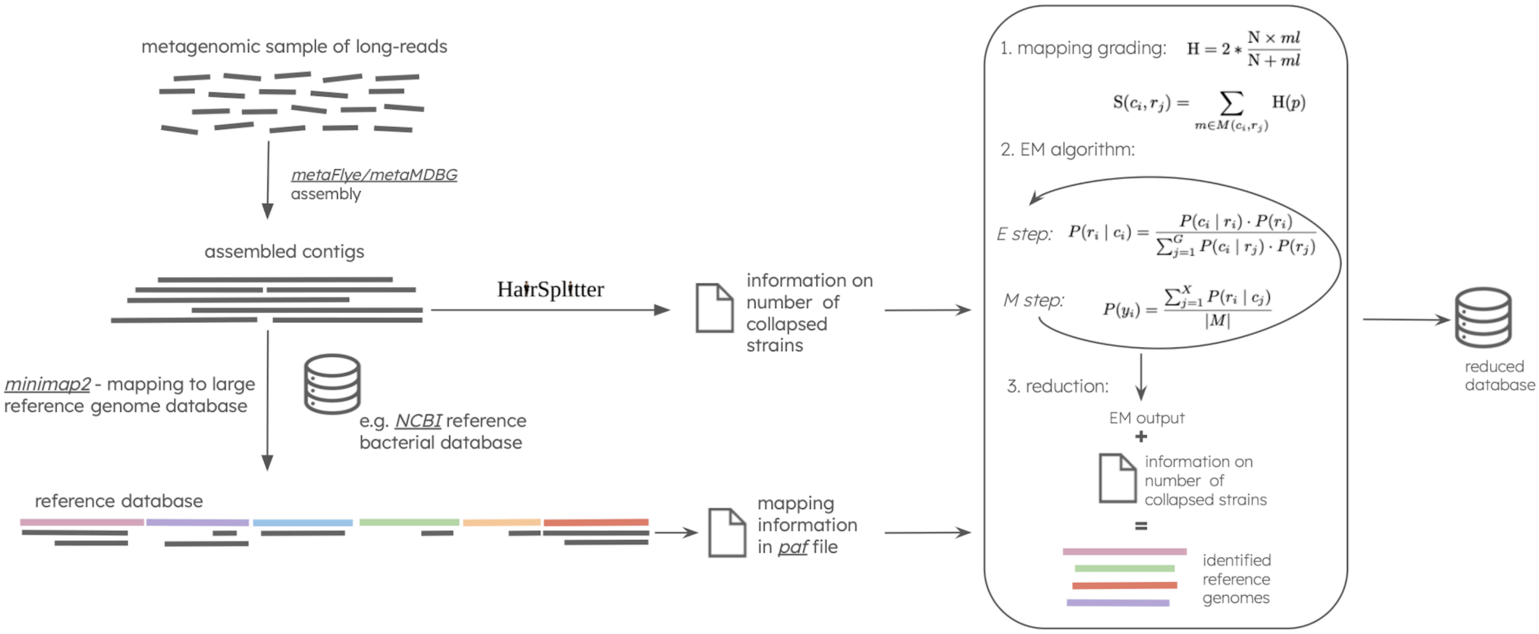


Figure S1: **MADRe Database Reduction pipeline.** The process begins with assembling long reads using metaFlye or metaMDBG. HairSplitter is then applied to estimate the number of collapsed strains per contig. The resulting contigs are mapped to the reference database, and the mapping information is stored in a PAF file. Organism identification within the reduced database is performed using an Expectation–Maximization (EM) algorithm that integrates both the mapping data and the estimated number of collapsed strains. The EM algorithm terminates after 25 iterations or when the change in pi values between two consecutive iterations (epsilon) falls below 0.0001. The number of collapsed strains estimated by HairSplitter is, by default, extended by two in MADRe, although this value can be adjusted through MADRe's parameters.

### Read Classification

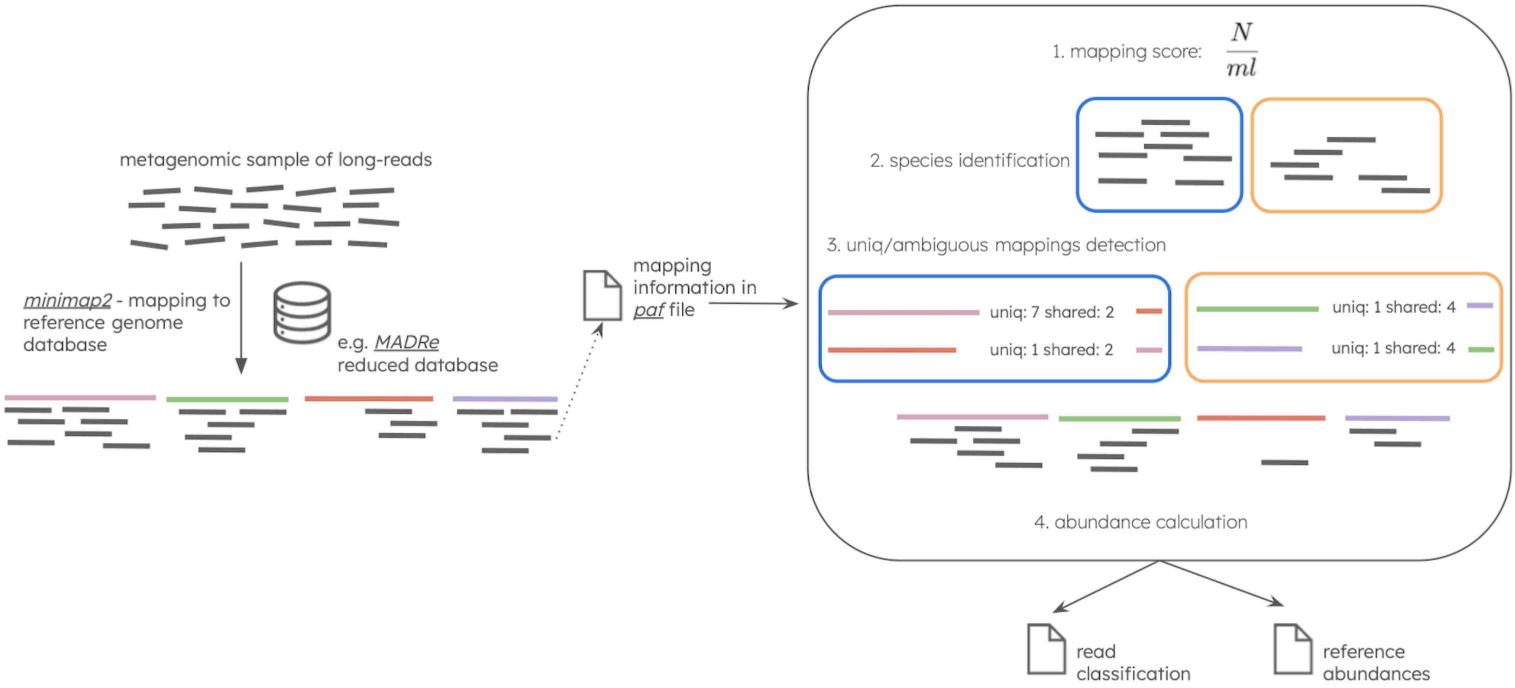


Figure S2: **MADRe Read Classification pipeline.** Reads are mapped to the (reduced) reference database, and the mapping information is stored in a PAF file. Based on mapping scores, reads are first grouped at the species level. Within each species, read assignments are further refined through a mapping-based probability reassignment procedure, which analyzes both unique and non-unique mapping profiles for each reference. In the final step, non-uniquely mapped reads are assigned to the reference with the highest number of unique and high-confidence non-unique mappings, as it is considered the most probable representative. Finally, reference-level abundances are computed.

### MADRe Clustering

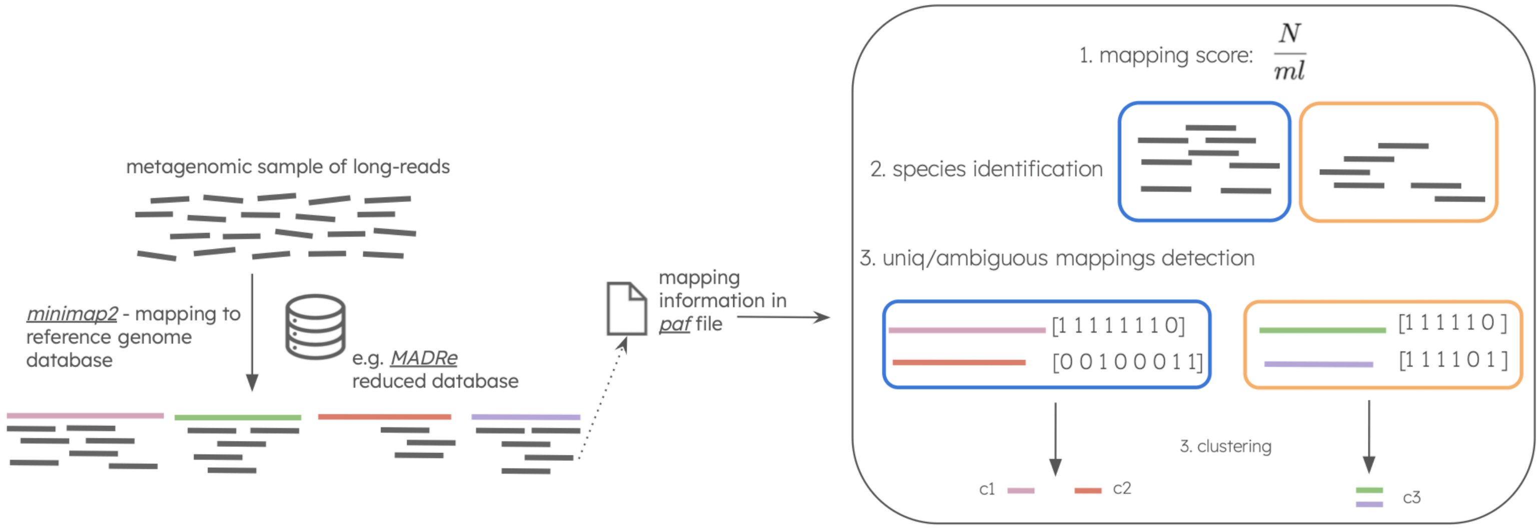


Figure S3: **MADRe Clustering pipeline.** Reads are mapped to the reference database, and the mapping information is stored in a PAF file. Based on mapping scores, reads are first grouped by species. For each species, every reference is represented by a binary vector, where each position corresponds to a read: a value of 1 indicates a high-quality mapping between the read and the reference, and 0 otherwise. These binary vectors are then used as input for DBSCAN clustering to group highly similar references (*eps*=0.9).

### Simulated data

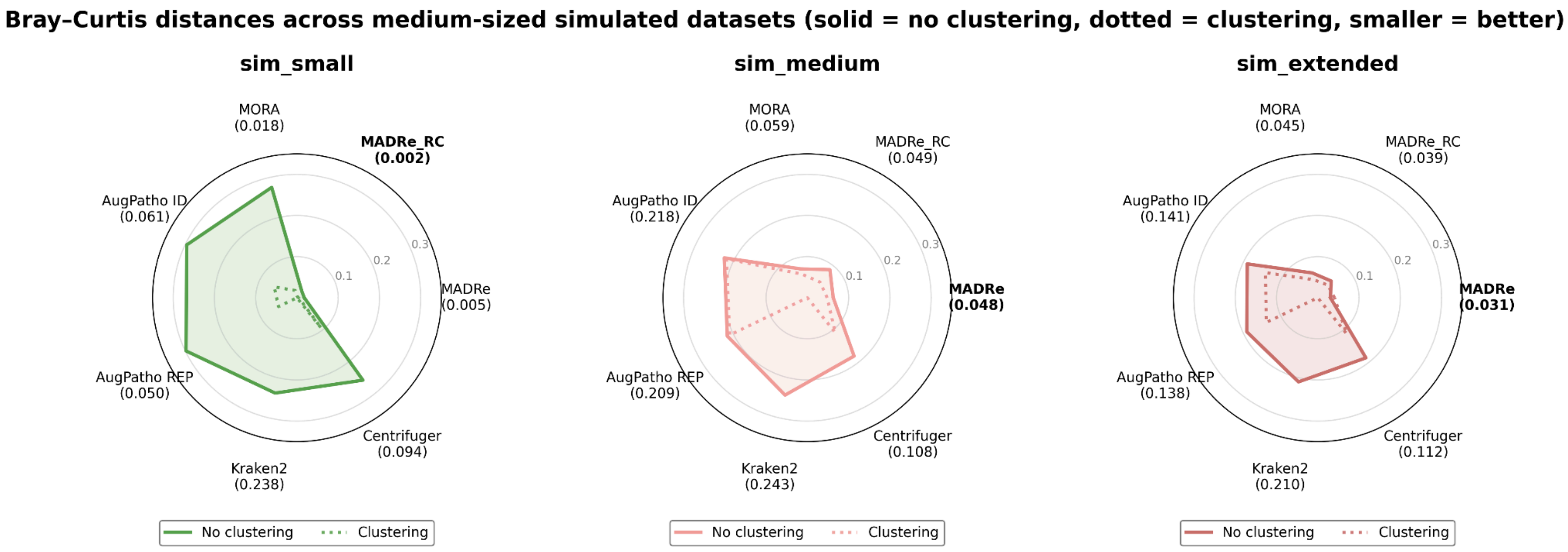


Figure S4: **Bray-Curtis distances for medium-sized simulated datasets.** The plots show BC distance between predicted and expected abundances in simulated datasets (sim_small, sim_medium, sim_extended) with and without post-clustering of similar strains. Since Kraken2 outputs taxID labels as classification labels, the clustering was not performed with it.

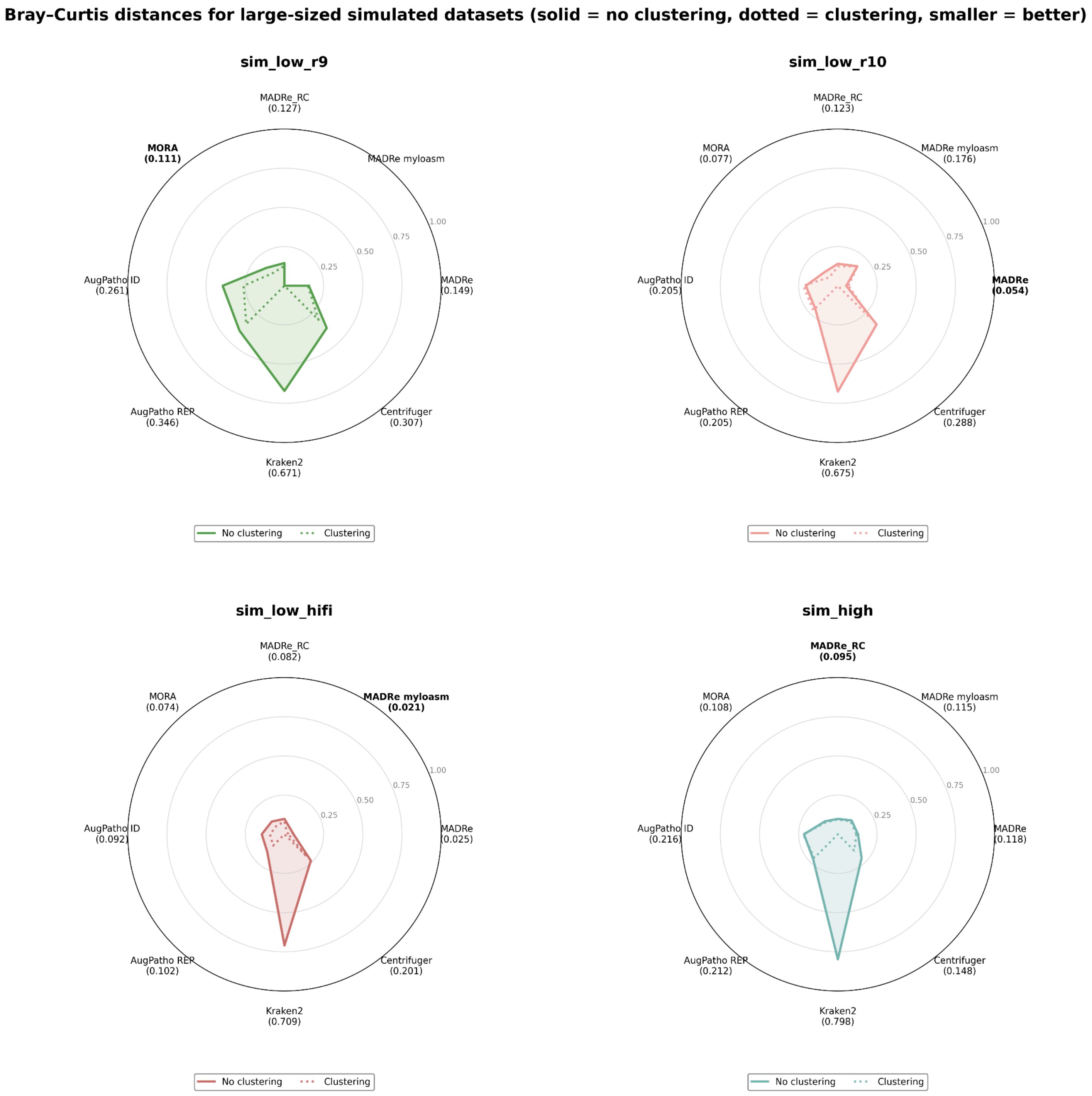


Figure S5: **Bray-Curtis distances for large-sized simulated datasets.** The plots show BC distance between predicted and expected abundances in simulated datasets (sim_low_r9, sim_low_r10, sim_low_hifi and sim_high) with and without post-clustering of similar strains. Since Kraken2 outputs taxID labels as classification labels, the clustering was not performed with it.

### Zymo data

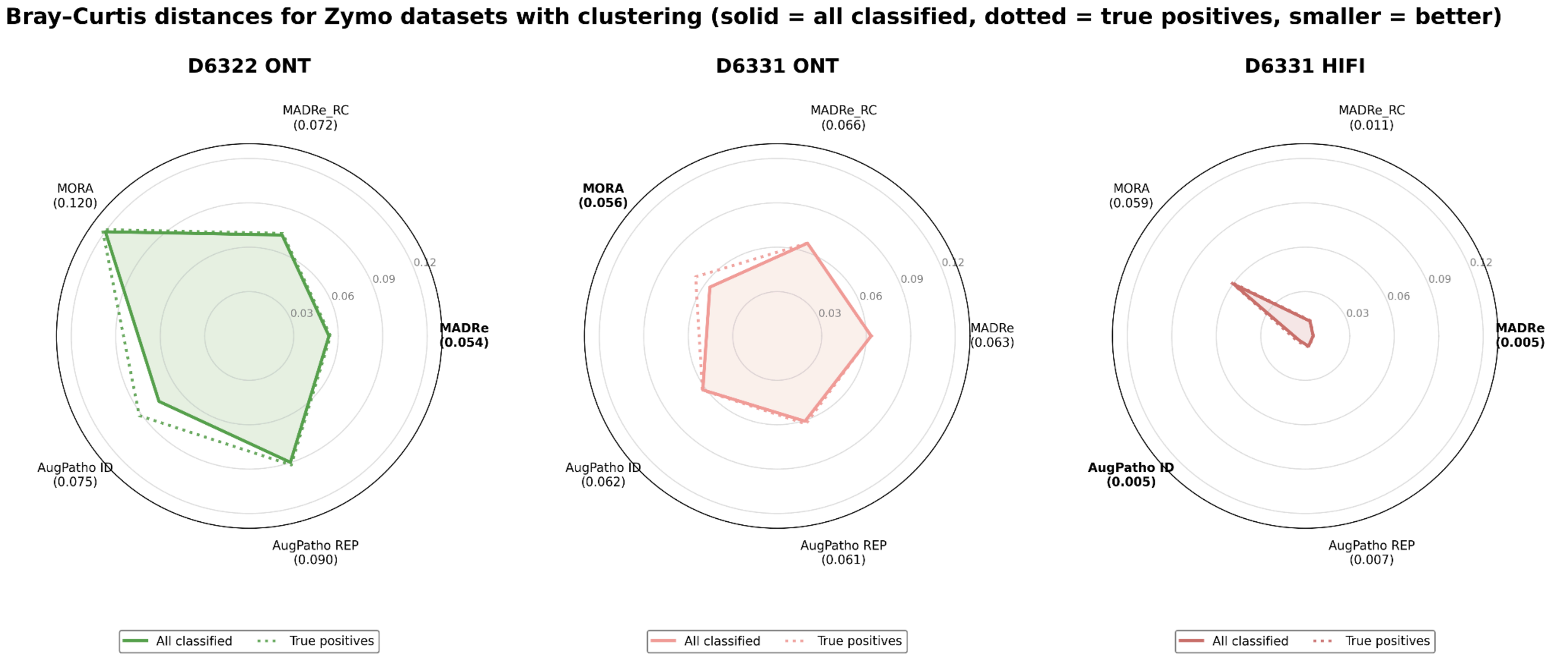


Figure S6: **Bray–Curtis distances for Zymo datasets with clustering of similar strains.** The plots show the BC distance between ground-truth read counts and classified read counts after post-classification clustering of similar strains. The solid line represents distances calculated using all classified reads, while the dotted line represents distances calculated using only true-positive classifications.

### Similar strains experiment

To investigate how MADRe behaves under specific conditions, we performed controlled experiments using six strains, including a trio of near-identical genomes (fastANI 99.98 - 99.9996%) (**Supplementary Table 23**). We compared MADRe with MORA and AugPatho (PathoScope2) - both purely mapping-based approaches - under four conditions: (i) all references present in the database, (ii) removal of closely related references from the database, (iii) altered strain abundances in the dataset, and (iv) removal of one strain from the dataset.

Across all scenarios, none of the tools proportionally distributed reads among the near-identical strains, instead, each displayed characteristic attractor behavior. MADRe consistently favored the centroid genome (1328|NZ_LR134283.1) - centroid according to ANI scores. AugPatho, which uses a penalized statistical mixture model for ambiguous reads, tended to collapse assignments onto a single dominant genome with slightly better alignment likelihoods, here 1671923|NZ_CP085939.1, even when its true abundance was low. MORA, which explicitly incorporates abundance constraints into its re-assignment optimization, showed context-dependent behavior: in some cases favoring 1671923|NZ_CP085939.1, in others over-allocating to absent genomes, reflecting the balance in its model between alignment scores and penalization for over-assignment.

When two references were removed (1328|NZ_LR134283.1 and 394340|NZ_AP024470.1), all methods reallocated reads primarily within the remaining near-identical genomes. MADRe maintained its centroid preference, AugPatho strongly collapsed reads of species 1328 into 1671923|NZ_CP085939.1, and MORA spread assignments more broadly but inflated 1328|NZ_AP018548.1 relative to truth. When abundances were changed (1671923|NZ_CP085939.1 was less abundant), MADRe and MORA under-recovered the reduced but nonzero 1671923|NZ_CP085939.1, while AugPatho continued to over-assign to 1671923|NZ_CP085939.1, consistent with its tendency to converge on a fixed attractor genome. Finally, in the last experiment when 1328|NZ_LR134283.1 was absent from the dataset, MADRe reassigned the majority of reads to 1671923|NZ_CP085939.1, which is the closest genome, despite 1328|NZ_LR134283.1 being the centroid when present. This shows that MADRe adapts to database composition. By contrast, AugPatho over-assigned to 1328|NZ_AP018548.1, and MORA produced a large spurious allocation to the absent 1328|NZ_LR134283.1 even though it is not in the dataset.

Overall, these results highlight a fundamental property of mapping-based tools: when an exact match is not available, each method employs its own strategy for forced classification. While AugPatho and MORA tend to collapse or redistribute ambiguous reads based on probabilistic or abundance-driven models, MADRe consistently seeks the most similar available genome (the centroid) ensuring that reads are assigned to the closest representative rather than arbitrarily redistributed.

### Real data

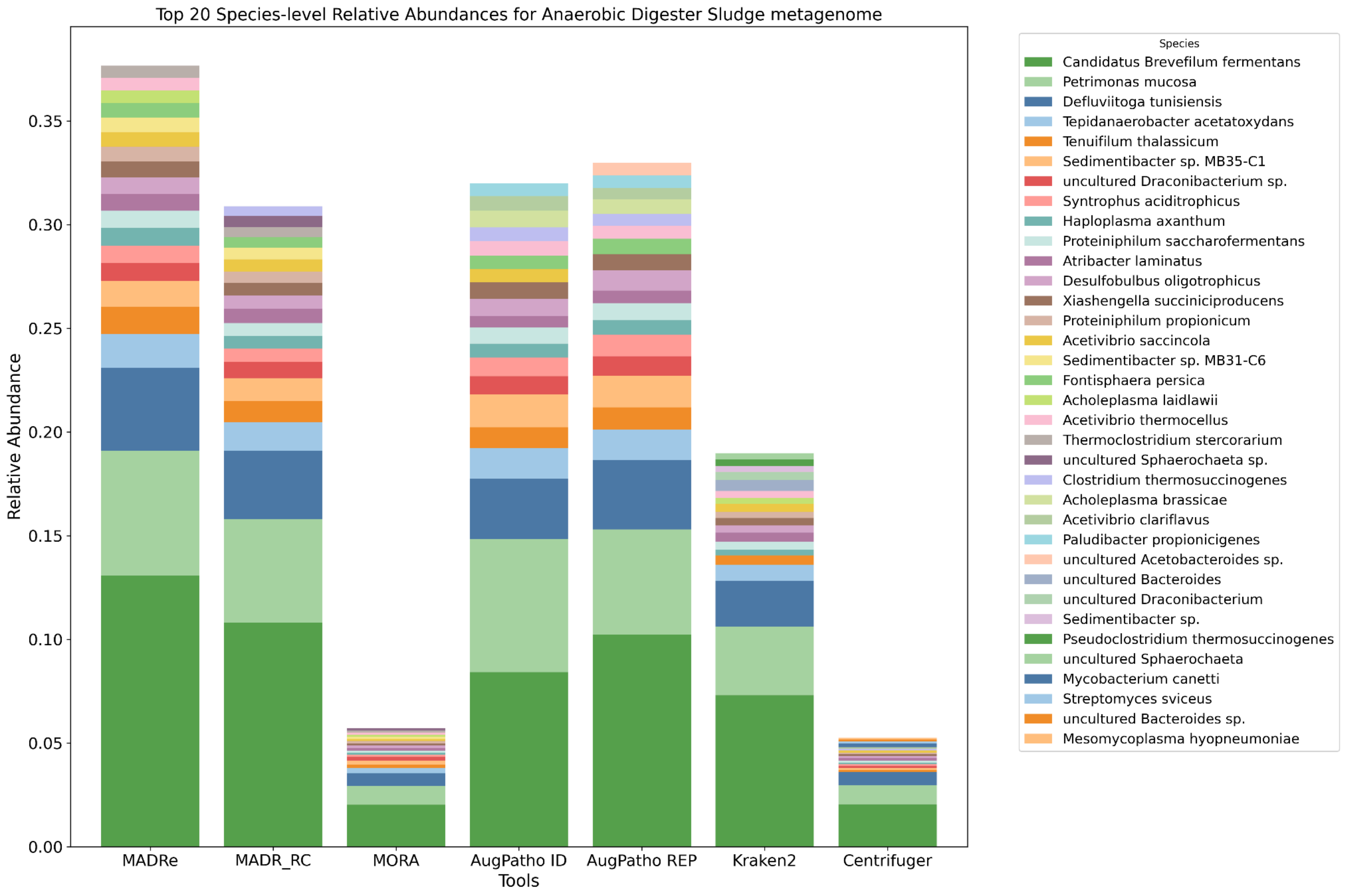


Figure S7: **Anaerobic Digester Sludge metagenome species-level abundances** of top-20 most abundant species for every of the tools. MORA's and Centrifuger's top 20 species account for the lowest cumulative abundance, suggesting its assignments are more broadly spread across a larger number of species.
